# Social networks predict the life and death of honey bees

**DOI:** 10.1101/2020.05.06.076943

**Authors:** Benjamin Wild, David M Dormagen, Adrian Zachariae, Michael L Smith, Kirsten S Traynor, Dirk Brockmann, Iain D Couzin, Tim Landgraf

**Author notes:** Equal contribution (Benjamin Wild, David M Dormagen, Adrian Zachariae).

## Abstract

In complex societies, individuals’ roles are reflected by interactions with other conspecifics. Honey bees (*Apis mellifera*) generally change tasks as they age, but developmental trajectories of individuals can vary drastically due to physiological and environmental factors. We introduce a succinct descriptor of an individual’s social network that can be obtained without interfering with the colony. This *network age* accurately predicts task allocation, survival, activity patterns, and future behavior. We analyze developmental trajectories of multiple cohorts of individuals in a natural setting and identify distinct developmental pathways and critical life changes. Our findings suggest a high stability in task allocation on an individual level. We show that our method is versatile and can extract different properties from social networks, opening up a broad range of future studies. Our approach highlights the relationship of social interactions and individual traits, and provides a scalable technique for understanding how complex social systems function.

## 1 Introduction

In complex systems, intricate global behaviors emerge from the dynamics of interacting parts. Within animal groups, studying the ways that individuals interact helps elucidate their function (Farine et al. 2015; Gordon 2010; Krause et al. 2015; Pinter-Wollman, Hobson, et al. 2014). Social interaction networks have been used to investigate e.g. pair-bonding (Psorakis et al. 2012), inter-group brokering (Lusseau et al. 2004), offspring survival (Cheney et al. 2016), cultural spread (Aplin et al. 2015), policing behavior (Flack et al. 2006), leadership in mobile groups (Strandburg-Peshkin et al. 2018), organization of food retrieval (Planckaert et al. 2019), and the propagation of alarm response (Rosenthal et al. 2015; Sosna et al. 2019). As our ability to collect detailed social network data increases, so too does our need to develop tools for understanding the significance and functional consequences of these networks (Gomez-Marin et al. 2014).

Social insects are an ideal model system to study the relationship between social interactions and individual roles, because task allocation has long been hypothesized to arise from interactions (Gordon 1996; Tofts et al. 1992; Traniello et al. 1997). The relationship of individual roles within the colony and the social network, however, is not well understood. Individuals, for example, can modify their behavior based on nestmate interaction (Pinter-Wollman, Bala, et al. 2013; Pinter-Wollman, Wollman, et al. 2011; Seeley 1992), and interactions change depending on where and with whom individuals interact (Davidson et al. 2017; Pinter-Wollman, Wollman, et al. 2011; Quevillon et al. 2015). These studies typically target specific types of interactions (e.g. food-exchange), specific roles within task allocation (e.g. foraging), or specific stimuli within the nest (e.g. brood), but an automatic observation system could capture the entirety of behaviors and interactions within a colony without human bias. Measuring the multitude of social interactions and their effect on behavior, and the social networks over the lifetime of individuals without interfering with the system (e.g., by removing individuals) is an open problem.

In honey bees, task allocation is characterized by temporal polyethism (Baracchi et al. 2014; Naug 2008; Schneider et al. 2004), where workers gradually change tasks as they age: young bees care for brood in the center of the nest, while old bees forage outside (Huang et al. 1996; Seeley 1982). Previous works often used few same-aged cohorts resulting in an unnatural age distribution (Baracchi et al. 2014; Gernat et al. 2018; Huang et al. 1996; Naug 2008). The developmental trajectory of individuals can, however, vary drastically due to internal physiological factors (i.e. genetics, ovary size, sucrose responsiveness (Gro Vang Amdam et al. 2003; Ihle et al. 2010; Pankiw et al. 1999; Scheiner 2012; Wang, Kocher, et al. 2012; Wang, Kaftanoglu, Siegel, et al. 2010)), nest state (i.e. amount of brood, brood age, food stores (Dreller et al. 1999; Seeley 1989; Traynor et al. 2015)), and the external environment (i.e. season, resource availability, forage success (Ament et al. 2010; Huang et al. 1995; Toth et al. 2005; Wang, Kaftanoglu, Brent, et al. 2016)). These myriad influences on maturation rate are difficult to disentangle, but all drive the individual’s behavior and task allocation. Due to the spatial organization of honey bee colonies, task changes also result in a change of location, with further implications on the cues that workers encounter (Seeley 1982). How and when bees change their allocated tasks in a natural setting has typically been assessed through destructive sampling (e.g. for measuring hormone titers of selected individuals), but understanding how all these factors combine would ideally be done in an undisturbed system.

With the advent of automated tracking, there has been renewed interest in how interactions change within colonies (Blut et al. 2017; Mersch et al. 2013), how spatial position predicts task allocation (Crall et al. 2018), and how spreading dynamics occur in social networks (Gernat et al. 2018). Despite extensive work on the social physiology of honey bee colonies (Seeley 1995), few works have studied interaction networks from a colony-wide or temporal perspective (Gernat et al. 2018; Hasenjager et al. 2020). While there is considerable variance in task allocation, even among bees of the same age, it is unknown to what extent this variation is reflected in the social networks. In large social groups, like honey bee colonies, typically only a subset of individuals are tracked, or tracking is limited to short time intervals (Blut et al. 2017; Bozek et al. 2020; Naug 2008).

Tracking an entire colony over a long time would allow one to investigate the stability of task allocation. Prior research has shown that during each life stage, an individual spends most of its time in a specific nest region (Johnson 2010; Seeley 1982), interacting with nestmates, but with whom they interact may depend on more than location alone (Farina 2000; Girard et al. 2011). Social interactions permit an exchange of information and can have long-term effects on an individual’s behavior (Cholé et al. 2019). While honey bees are well-known for their elaborate social signals (e.g. waggle dance, shaking signal, stop signal (Frisch 1967; Nieh 2010; Seeley et al. 2012)), they also exchange information through food exchange, antennation, or simple colocalization (Balbuena et al. 2012; Goyret et al. 2003). However, identifying an individual’s role in a group based on its characteristic patterns of interaction remains challenging, particularly with large numbers of individuals and multiple modes of interaction.

In order to broadly understand how an individual’s social network may contribute to its lifetime role within a complex society, we developed a tracking method of an entire colony of several thousand individuals with a natural age distribution, which permits unbiased long-term assessments. We investigate how we can extract meaning from the multitude of interactions and interaction types, among thousands of individuals in a honey bee colony.

We introduce a low-dimensional descriptor, network age, that allows us to compress the social network of all individuals in the colony into a single number per bee per day. Network age, and therefore the social network of a bee, captures the individual’s behavior and social role in the colony and allows us to predict task allocation, mortality, and behavioral patterns such as velocity and circadian rhythms. Following the developmental trajectories of individual honey bees and cohorts that emerged on the same day reveals clusters of different developmental paths, and critical transition points. In contrast to these distinct clusters of long-term trajectories, we find that transitions in task allocation are fluid on an individual level. We show that the task allocation of individuals in a natural setting is stable over long periods, allowing us to predict a worker’s task better than biological age up to one week into the future.

## 2 What is network age?

To obtain the social network structure over the lifetime of thousands of bees, we require methods that will track the tasks and social interactions of many individuals over consecutive days. We video recorded a full colony of individually marked honey bees (*A. mellifera*) at 3 Hz for 25 days (from 2016-08-01 to 2016-08-25) and obtained continuous trajectories for all individuals in the hive (Boenisch et al. 2018; Wario, Wild, Couvillon, et al. 2015). We used a two-sided single-frame observation hive with a tagged queen and started introducing individually tagged bees into the colony approximately one month before the beginning of the focal period (2016-06-28 to 2016-08-23). To ensure that no unmarked individuals emerged inside the hive, we replaced the nest substrate regularly (approx. every 21 days). In total, we recorded 1921 individuals aged from 0 days to 8 weeks.

A worker’s task and the proportion of time she spends in specific nest areas are tightly coupled in honey bees (Seeley 1982). We annotated nest areas associated with specific tasks (e.g. brood area or honey storage) for each day separately (see section SI 1.4), as they can vary in size and location over time (Smith et al. 2016). We then use the proportion of time an individual spends in these areas throughout a day as an estimate of her current tasks.

We calculated daily aggregated interaction networks from contact frequency, food exchange (trophallaxis), distance, and changes in movement speed after contacts (see section SI 1.3). These networks contain all the pairwise interactions between all individuals over time. For each day and interaction type, we extract a compact representation that groups bees together with similar interaction patterns, using spectral decomposition (Belkin et al. 2003; von Luxburg 2007). We then combine each bees’ daily representations of all interaction types and map them to a scalar value (network age) that best reflects the fraction of time spent in the task-associated areas using CCA (canonical-correlation analysis; (Hotelling 1936; Knapp 1978)). Note that network age is solely a representation of the social network and not of location; the fraction of time spent in the task-associated areas is only used to select which information to extract from the social networks (e.g., by assigning higher importance to trophallactic interactions). Network age can still represent an individual’s location, but only if this information is inherently present in the social networks. Network age thus compresses millions of data points per individual and day (1920 potential interaction partners, each detected 127 501 ± 50 340 times on average per day, with four different interaction types) into a single number that represents each bee’s daily position in the multimodal temporal interaction network. Since CCA is applied over the entire focal period, network age can only represent interaction patterns that are consistent over time. See fig. 1 for an overview and section SI 1.5 for a detailed description of the methods.

**Figure 1:**
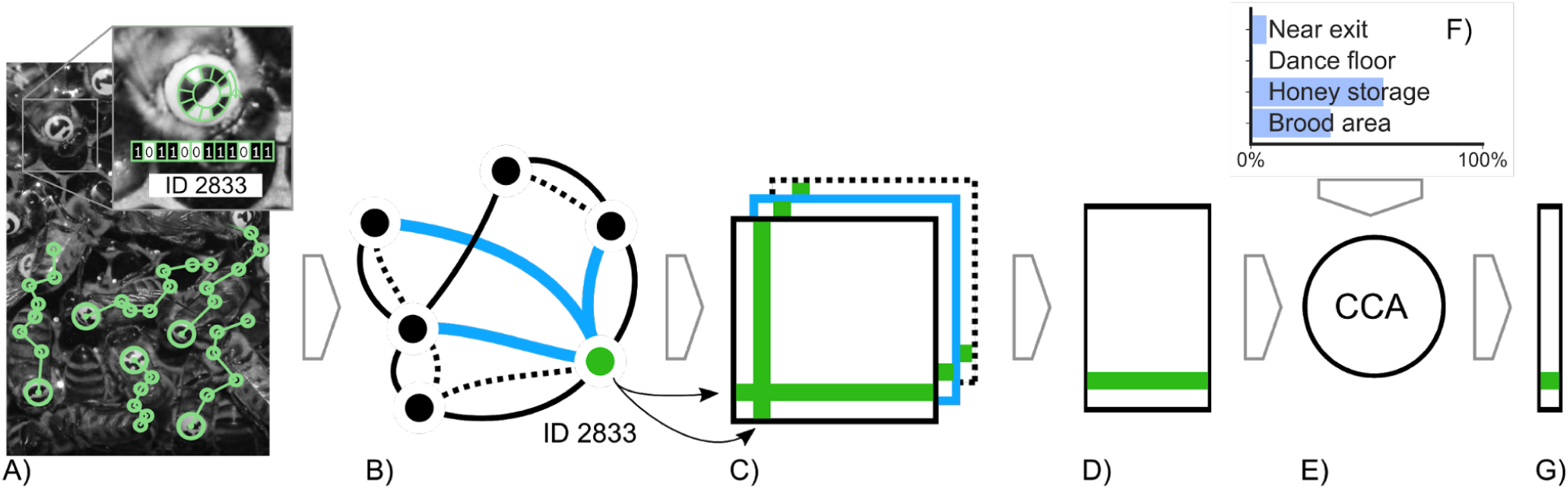
Network age, a one-dimensional descriptor of an individual’s role within the colony, based on an individual’s interaction pattern. Using the BeesBook automated tracking system, we obtain lifetime tracking data for individuals (A). These tracks are used to construct multiple weighted social interaction networks (B). We aggregate daily networks (C) to then extract embeddings that group bees together with similar interaction patterns, using spectral decomposition (D). Finally, we use a linear transformation (e; CCA = canonical-correlation analysis) that maximizes correlation with the fraction of time spent in different nest areas (F) to compress them into a single number called *network age* (G).

Network age is a unitless descriptor. We scale it such that 90% of the values are between 0 and 40 to make it intuitively comparable to a typical lifespan of a worker bee during summer, and because biological age is commonly associated with task allocation in honey bees. This scaling can be omitted for systems where behavior is not coupled with biological age.

## 3 Network age correctly identifies task allocation

We quantify to what extent network age, and therefore the social interaction patterns, reflect task allocation by using multinomial regression to predict the fraction of time each bee spends in the annotated nest areas (see section SI 2.4). Note that while we also used these spatial preferences to select which information to extract from the interaction networks, it is not certain whether the spatial information is contained in the social network in the first place, and how well a single dimension can capture it. Furthermore, the social network structure could vary over many days with changing environmental influences, preventing the extraction of a stable descriptor. The regression analysis allows us to compare different variants of network age to biological age as a reference.

To evaluate the regression fit, we use McFadden’s pseudo *R*^2^ scores 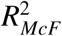 (McFadden 1973). Network age is twice as good as biological age at predicting the individuals’ location preferences, and therefore their tasks (network age: median 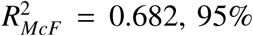 CI [0.678, 0.687]; biological age: median 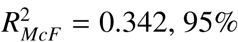 CI [0.335, 0.349]; 95% CI of effect size [0.332, 0.348], N=128; likelihood-ratio *χ*^2^ test *p* ≪ 0.001, N=26 403, table 1, see section SI 2.6 for details). Network age provides a better separability of time spent in task-associated nest areas than biological age (fig. 2A, example cohort in fig. 2C, section SI 1.8 for all cohorts). Network age correlates with location because of the inherent coupling between tasks and nest areas. Still, it is not a direct measure of location: bees with the same network age can exhibit different spatial distributions and need not directly interact (see section SI 2.3).

**Table 1:** 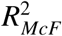 median scores and bootstrapped 95% CI for different combinations of independent variables in the rows and dependent variables in the columns. We evaluated the following independent variables: different dimensionalities of network age (*Network age, Network age (2D), Network age (3D)*, see section SI 1.5), network age derived without supervision (*PCA*, see section SI 1.6), with a subsample of the bees (*1% of bees tagged* and *5% of bees tagged*, see section SI 2.2), and based only on spatial proximity (*Euclidean proximities* and *Proximity events*). The likelihood ratio test was evaluated for the combined models (e.g. *Age* + *Network age*, see section SI 2.6).

|  | Brood area | Dance floor | Honey storage | Near exit | Combined |
| --- | --- | --- | --- | --- | --- |
| Age | 0.46 [0.45, 0.47] | 0.28 [0.27, 0.29] | 0.00 [0.00, 0.00] | 0.32 [0.31, 0.33] | 0.34 [0.34, 0.35] |
| Age (nonlinear) | 0.51 [0.50, 0.52] | 0.35 [0.34, 0.36] | 0.09 [0.09, 0.10] | 0.41 [0.40, 0.42] | 0.38 [0.38, 0.39] |
| Network age (1% of bees tagged) | 0.58 [0.07, 0.79] | 0.50 [0.08, 0.68] | 0.03 [0.00, 0.21] | 0.49 [0.12, 0.73] | 0.52 [0.13, 0.70] |
| Network age (Only euclidean proximities) | 0.51 [0.50, 0.52] | 0.37 [0.36, 0.38] | 0.17 [0.16, 0.18] | 0.53 [0.52, 0.54] | 0.52 [0.51, 0.52] |
| Network age (Only euclidean proximities, nonlinear) | 0.60 [0.60, 0.61] | 0.55 [0.54, 0.56] | 0.20 [0.18, 0.21] | 0.56 [0.55, 0.57] | 0.55 [0.55, 0.56] |
| Network age (1% of bees tagged, nonlinear) | 0.62 [0.08, 0.81] | 0.58 [0.16, 0.73] | 0.16 [0.03, 0.34] | 0.55 [0.17, 0.77] | 0.55 [0.19, 0.72] |
| Network age (5% of bees tagged) | 0.76 [0.66, 0.83] | 0.61 [0.52, 0.68] | 0.03 [0.00, 0.08] | 0.60 [0.52, 0.69] | 0.65 [0.58, 0.70] |
| Network age (PCA) | 0.80 [0.79, 0.80] | 0.53 [0.52, 0.54] | 0.00 [0.00, 0.00] | 0.58 [0.57, 0.59] | 0.65 [0.64, 0.65] |
| Network age (PCA, nonlinear) | 0.80 [0.80, 0.80] | 0.61 [0.60, 0.62] | 0.27 [0.26, 0.28] | 0.66 [0.65, 0.66] | 0.66 [0.66, 0.67] |
| Network age (Only proximity events) | 0.79 [0.79, 0.80] | 0.64 [0.64, 0.65] | 0.03 [0.03, 0.03] | 0.60 [0.59, 0.60] | 0.67 [0.67, 0.68] |
| Network age (5% of bees tagged, nonlinear) | 0.77 [0.63, 0.83] | 0.66 [0.58, 0.73] | 0.24 [0.15, 0.32] | 0.67 [0.57, 0.73] | 0.67 [0.59, 0.72] |
| Network age | 0.80 [0.80, 0.81] | 0.63 [0.63, 0.64] | 0.03 [0.03, 0.04] | 0.61 [0.60, 0.62] | 0.68 [0.68, 0.69] |
| Age + Network age | 0.81 [0.80, 0.81] | 0.63 [0.63, 0.64] | 0.10 [0.09, 0.11] | 0.61 [0.60, 0.62] | 0.69 [0.69, 0.69] |
| Network age (Only proximity events, nonlinear) | 0.81 [0.80, 0.81] | 0.68 [0.67, 0.69] | 0.32 [0.31, 0.32] | 0.68 [0.67, 0.69] | 0.70 [0.69, 0.70] |
| Network age (nonlinear) | 0.82 [0.81, 0.82] | 0.68 [0.67, 0.69] | 0.33 [0.32, 0.33] | 0.69 [0.69, 0.70] | 0.71 [0.70, 0.71] |
| Age + Network age (nonlinear) | 0.82 [0.82, 0.83] | 0.69 [0.68, 0.69] | 0.36 [0.35, 0.37] | 0.70 [0.69, 0.70] | 0.72 [0.71, 0.72] |
| Network age (2D) | 0.84 [0.84, 0.85] | 0.65 [0.64, 0.65] | 0.46 [0.45, 0.46] | 0.64 [0.63, 0.64] | 0.73 [0.73, 0.74] |
| Network age (2D, nonlinear) | 0.85 [0.85, 0.85] | 0.69 [0.69, 0.70] | 0.49 [0.48, 0.50] | 0.70 [0.70, 0.71] | 0.74 [0.74, 0.75] |
| Network age (3D) | 0.84 [0.84, 0.85] | 0.71 [0.70, 0.71] | 0.46 [0.45, 0.47] | 0.70 [0.70, 0.71] | 0.76 [0.75, 0.76] |
| Network age (3D, nonlinear) | 0.86 [0.85, 0.86] | 0.75 [0.74, 0.75] | 0.50 [0.49, 0.51] | 0.76 [0.76, 0.77] | 0.77 [0.77, 0.77] |

**Figure 2:**
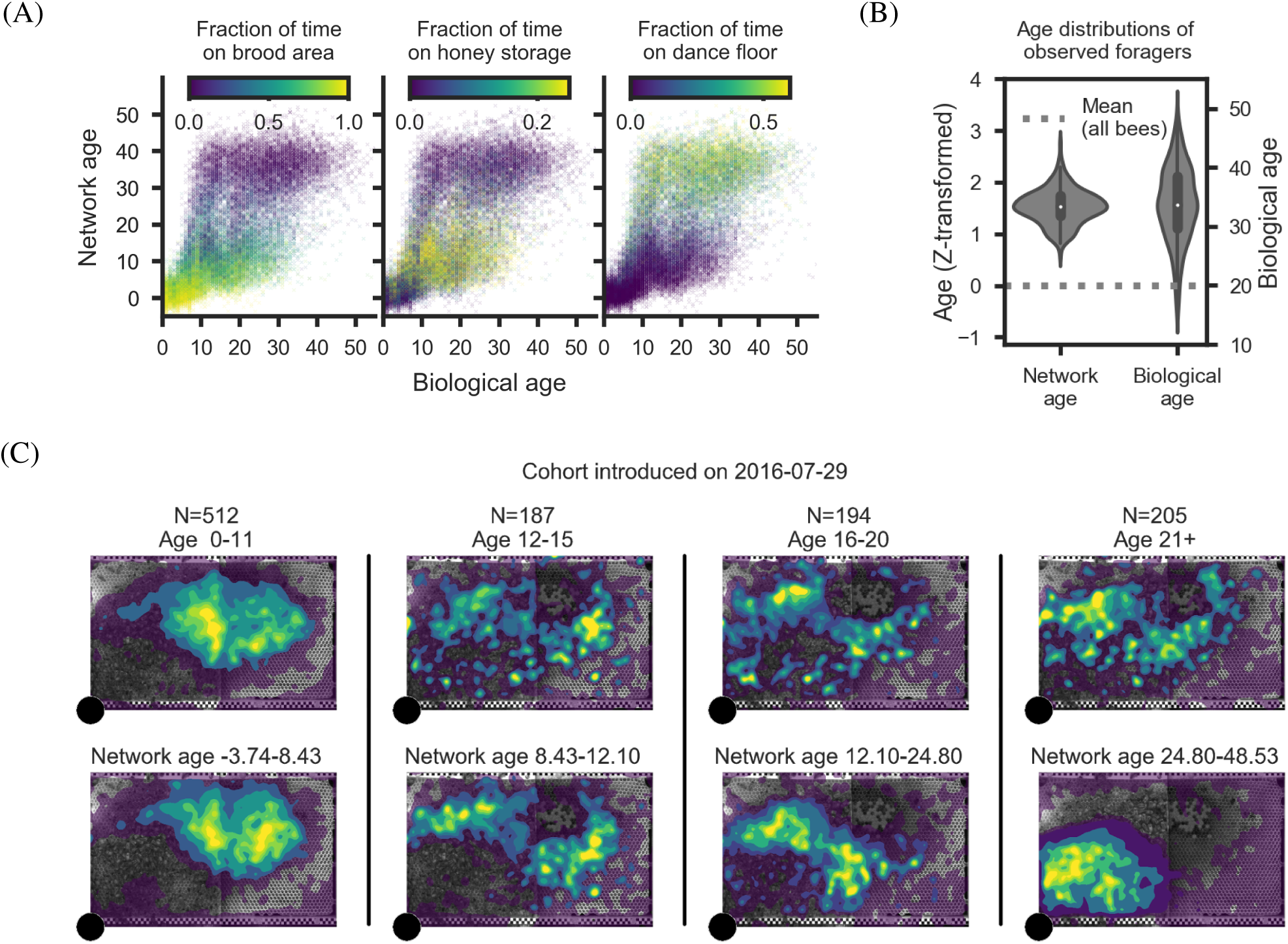
Network age is an accurate descriptor of task allocation. (A) The proportion of time spent on task-associated locations in relation to biological age and network age with each cross representing one individual on one day of her life. For a given value on the y-axis (network age) colors are more consistent than for a given value on the x-axis (biological age). (B) Z-transformed age distributions for known foragers visiting a feeder (N=40). The variance in biological age is greater than the variance in network age (boxes: center dot, median; box limits, upper and lower quartiles; whiskers, 1.5x interquartile range). Corresponding biological ages are also shown on the right y-axis (original biological age: 34.2 ± 7.9, original network age: 38.3 ± 4.6; mean standard deviation). (C) Spatial distributions of an example cohort over time (bees emerged on 2016-07-29, 64 individuals over 25 days), grouped by biological age (top row) versus network age (bottom row). Note how network age more clearly delineates groups of bees than biological age, with bees transitioning from the brood nest (center of the comb), to the surrounding area, to the dance floor (lower left area).

While we can improve the predictive power of network age by extracting a multi-dimensional descriptor instead of a single value (see sections SI 1.5 and SI 2.4 for details), the improvements for additional dimensions are marginal. This implies that a one-dimensional descriptor captures most of the information from the social networks that is relevant to the individuals’ location preferences and therefore their tasks.

We experimentally demonstrated that network age robustly captures an individual’s task by setting up sucrose feeders and identifying workers that foraged at the feeders (known foragers, N=40, methods in section SI 2.5). We then compared the biological ages of these known foragers to their network ages. As expected, foragers exhibited a high biological age and a high network age, whereas biological age exhibited significantly larger variance than network age (fig. 2B; Levene’s test (Levene 1960), performed on z-transformed values: *p* ≪ 0.001, N=200). Indeed, while we observed a forager as young as 12 days old, that individual had a network age of 25.5, demonstrating that network age more accurately reflected her task than her biological age (z-transformed values: biological age −0.46; network age 0.61).

Tagging an entire honey bee colony is laborious. However, by sampling subsets of bees, we find that network age is still a viable metric, even when only a small proportion of individuals are tagged and tracked. With only 1% of the bees tracked, network age is still a good predictor of task (median 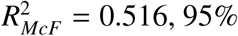 CI [0.135, 0.705], N=128) while increasing the number of tracked individuals to 5% of the colony results in a 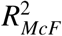 value comparable to the fully tracked colony (5% of colony tracked: median 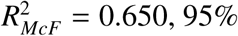 CI [0.578, 0.705], N=128; whole colony tracked: median 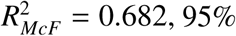 CI [0.678, 0.687], N=128; see section SI 2.2). Similarly, we find that an approximation of network age can be calculated without annotated nest areas: Network age can be extracted in an unsupervised manner using PCA on the spectral embeddings of the different interaction type matrices (median 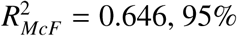 CI [0.641, 0.650], N=128, see section SI 1.6).

## 4 Developmental changes over the life of a bee

Network age reveals differences in interaction patterns and task allocation among same-aged bees (fig. 3A). After around six days of biological age, the network age distribution becomes bimodal (see section SI 3.4). Bees in the functionally old group (high network age) spend the majority of their time on the dance floor, whereas same-aged bees in the functionally young group (low network age) are found predominantly in the honey storage area (fig. 3A). Transitions from high to low network age are a rare occurrence in our colony (see section SI 3.5).

**Figure 3:**
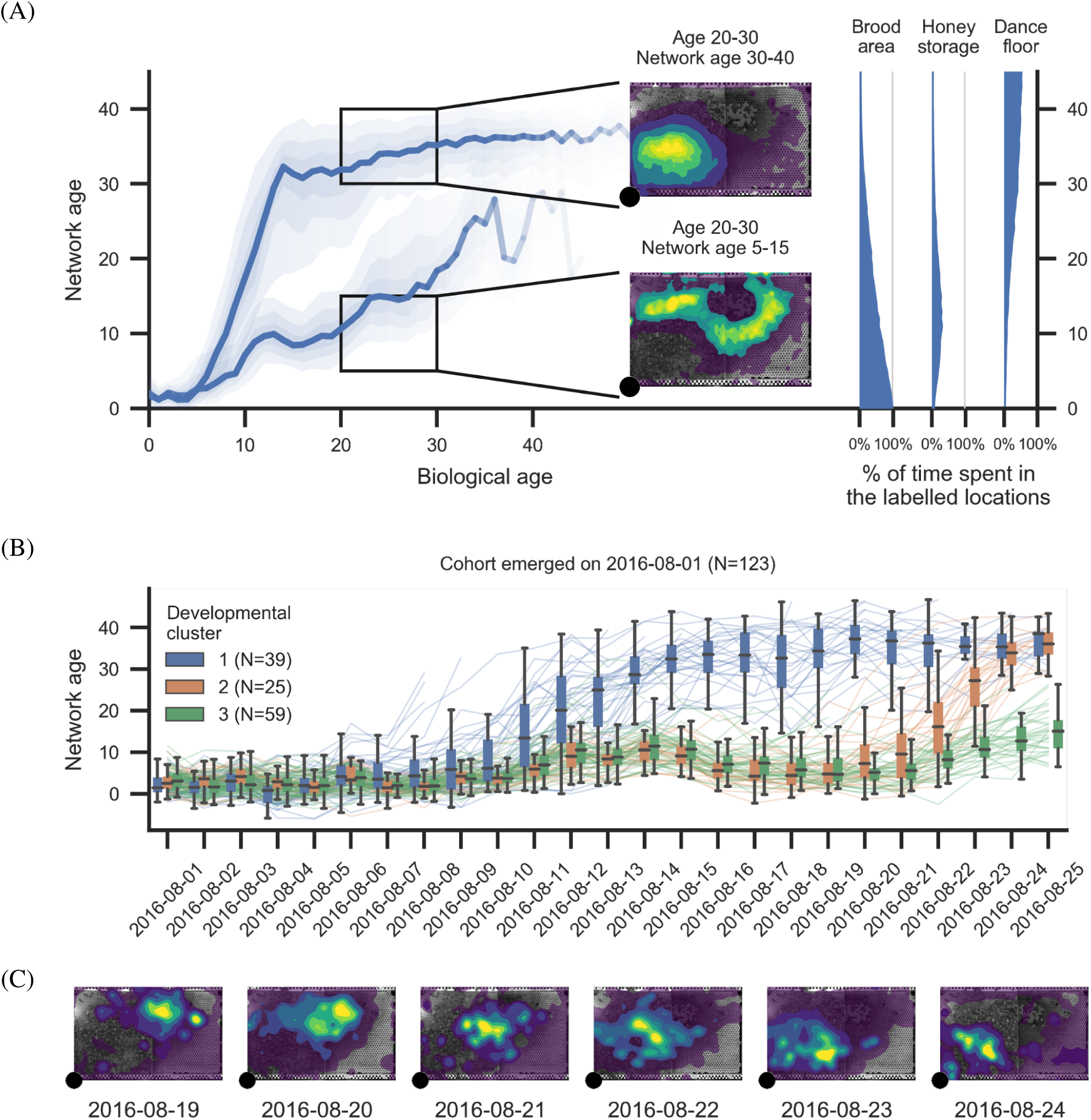
Network age reveals distinct developmental paths. (A) Left: The median of network ages over biological ages for all individuals that lived more than 11 days (light blue: 60% of data; see section SI 3.3). We observe a split in network age corresponding to different tasks: The upper heatmap (network age 30-40, biological age 20-30, 577 bees, 857 283 data points) corresponds to the dance floor, while the lower heatmap (network age 5-15, biological age 20-30, 381 bees, 742 622 data points) borders between dance floor and brood nest. Right: The mean fraction of time a bee with a given network age spends on our annotated regions throughout a day. (B) Lines depict the network age of individual bees of a same-aged cohort with the colors indicating clusters of their network age over time. Boxes summarize bees belonging to each cluster for a given day (center line, median; box limits, upper and lower quartiles; whiskers, 1.5x interquartile range). (C) Heatmaps showing the spatial distribution of bees in the developmental cluster 2 (orange) from 2016-08-19 to 2016-08-24. The smooth transition in network age (orange in line plot, B) from one mode to another corresponds to a smooth transition in spatial location (heatmaps, C).

We attribute the split on the population level to distinct patterns of individual development. Clustering the time series of network ages over the lives of bees identifies distinct developmental paths within same-aged cohorts (see section SI 3.2 for the method). In the cohort that emerged on 2016-08-01, the first developmental cluster (blue, fig. 3B) rapidly transitions to a high network age (likely corresponding to foraging behavior) after only 11 days. The second cluster (orange) transitions at around 21 days of biological age, while bees in the third cluster (green) remain at a lower network age throughout the focal period. We see similar splits in developmental trajectories for all cohorts, although the timing of these transitions varies (see section SI 3.2 for additional cohorts). Such divergence in task allocation has been previously shown in bees; factors that accelerate a precocious transition to foraging include hormone titers (Robinson et al. 1987), genotype (Pankiw et al. 1999), physiology, especially the number of ovarioles (Gro V. Amdam, Norberg, et al. 2004), and sucrose response threshold (Scheiner et al. 2004).

The transition from low to high network age over multiple days is characterized by a gradual shift in the spatial distribution (see example in fig. 3C), highlighting that an individual’s task changes gradually. The network age of most bees is highly repeatable (median *R* = 0.612 95% CI=[0.199, 0.982], see section SI 3.1 for details), indicating task stability over multiple days. Both findings (gradual change over a few days and high repeatability) are consistent with the dynamics of the underlying physiological processes, such as vitellogenin and juvenile hormone, that influence task allocation and the transition to foraging (Gro V. Amdam and Page Jr 2010).

## 5 Network age predicts an individual’s behavior and future role in the colony

Network age predicts task allocation (i.e. in what part of the nest individuals will be) up to ten days into the future. Knowing the network age of a bee today allows a better prediction of the task performed by that individual next week than her biological age informs about her current tasks (fig. 4C, binomial test, *p* ≪ 0.001, N=55 390, 95% CI of effect size [0.055, 0.090], N=128, see section SI 4.4 for details). We confirm that this is only partially due to network age being repeatable (see section SI 4.4). We do note, however, that our ability to predict the future tasks of a young bee is limited, especially before cohorts split into high and low network age groups (fig. 3A). This limitation hints at a critical developmental transition point in their lives, an attractive area for future study.

**Figure 4:**
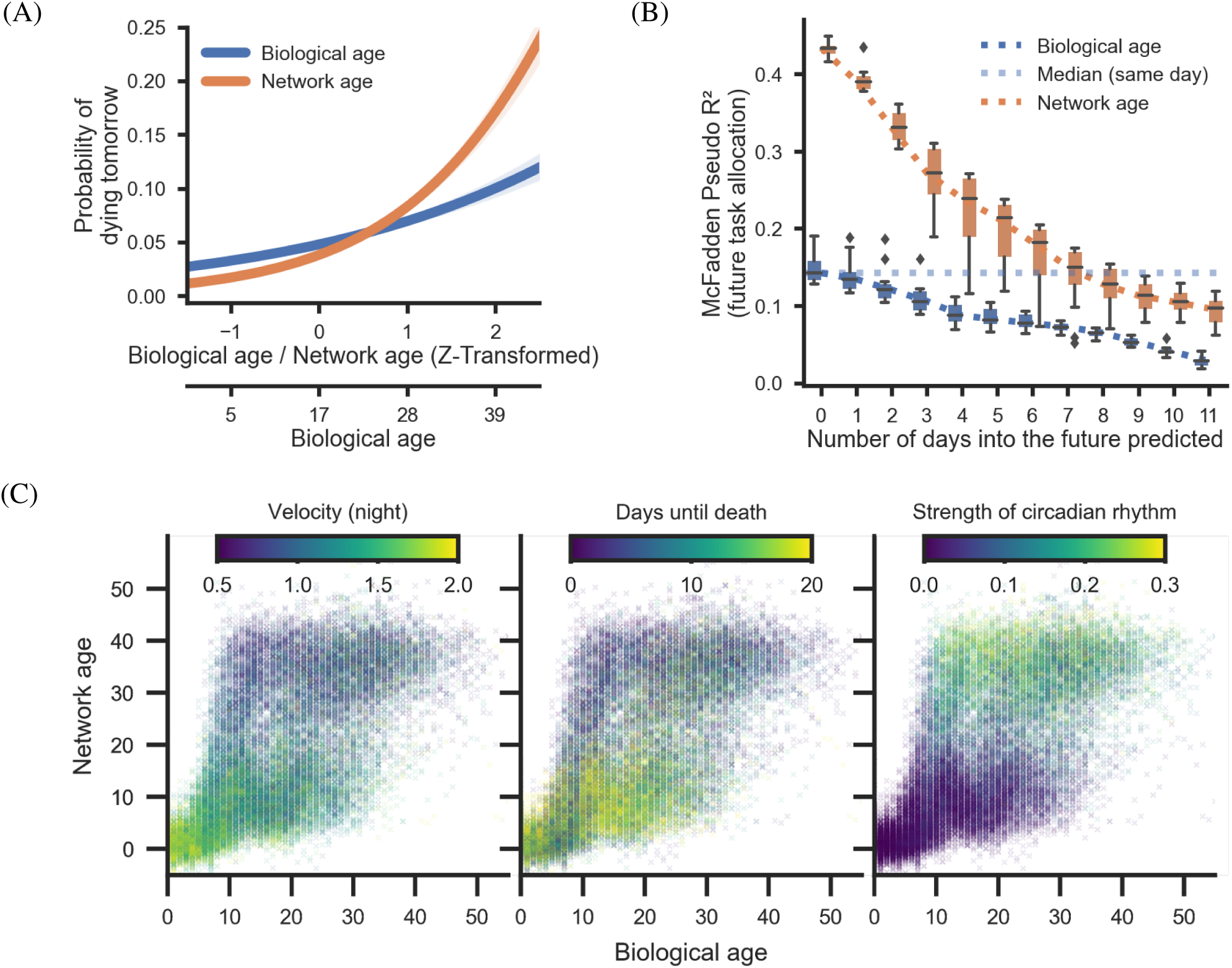
Network age can be used to predict other properties, such as mortality and circadian rhythms. It also predicts an individual’s future task allocation. (A) An individual’s mortality on the next day based on her age (x-axes show original and z-transformed biological age and z-transformed network age). Bees with a low network age have lower mortality than biologically young bees; bees with a high network age have higher mortality than biologically old bees (shaded areas: 95% bootstrap confidence intervals for the regression estimates). (B) Network age can be used to predict task allocation and future behaviors. Network age predicts the task of an individual 7 days into the future better than biological age predicts the individual’s task the same day (blue dotted line). Each box comprises 128 bootstrap samples (center line, median; box limits, upper and lower quartiles; whiskers, 1.5x interquartile range; points, outliers). (C) Selected properties mapped for network age over biological age with each cross representing one individual on one day of her life. Note that for a given value on the y-axis (network age) colors are more consistent than for a given value on the x-axis (biological age).

We explicitly optimized network age to be a good predictor of task-associated locations. However, we find that network age predicts other behaviors better than biological age, including an individual’s impending death (network age: median *R*^2^ = 0.165, 95% CI [0.158, 0.172], versus biological age: median *R*^2^ = 0:064, 95% CI [0.059, 0.068]; 95% CI of effect size [0.037, 0.039], N=128, likelihood-ratio *χ*^2^ test *p* ≪ 0.001, N=26 403). Biologically-young but network-old bees have a significantly higher probability of dying within a week (80.6% N=139) than do biologically-old but network-young bees (42.1% N=390; *χ*^2^ test of independence *p* ≪ 0.001 N=529; see section SI 4.5 for details). This is likely because a biologically-young bee with a high network age, i.e. a bee that starts to forage earlier in life and faces more perils imposed by the outside world, is more likely to die than a bee of the same age with a low network age. This finding is consistent with previous work showing increased mortality with precocious foraging (Perry et al. 2015; Rueppell et al. 2008).

Network age also captures movement patterns of individual bees such as daily and nightly average speed, the circadian rhythm, and the time of an individual’s peak activity better than biological age (for all dependent variables likelihood-ratio *χ*^2^ test *p* ≪ 0.001, N=26 403, see section SI 4.3 for 95% CI of effect sizes). Network age thus explains behaviors of individuals independent of their location.

To investigate whether network age is a good predictor of future task allocation and behavior only because it captures the spatial information contained in the social network, we repeat the analyses above using the time spent in task-associated locations as independent variables. We find that network age, even though it was extracted using this spatial information as a guide, is still a better predictor of an individuals’ behavior (for all dependent variables likelihood-ratio *χ*^2^ test *p* ≪ 0.001, N=26 403, see Location (1D) in section SI 4.6 and table 4d for 95% CI of effect sizes). This difference in predictive power highlights that the multimodal interaction network contains more information about an individual than spatial information alone.

While we focus on predicting tasks from network age, we can control the information we extract from the observed social networks and derive variants of network age better suited for other research questions. By replacing the *task associated location preferences* in the final step of our method with *days until death*, we extracted a descriptor that captures social interaction patterns related to mortality. This descriptor improves the prediction of the individuals’ death dates by 31% compared to network age (median increase in *R*^2^ = 0.05; 95% CI [0.04, 0.06] N=128, see section SI 4.2 and table 4c), opening up novel social network perspectives for studies such as the risk factors of disease transmission. Similarly, we extracted descriptors optimized to predict the movement patterns introduced in the last paragraph (for all except *Time of peak activity* likelihood-ratio *χ*^2^ test *p* ≪ 0.001, N=26 403, see table 4c for 95% CI of effect sizes). These targeted embeddings provide precise control over the type of information we extract from the social networks and extend the network age method to address other important research questions in honey bees and other complex animal societies.

## 6 Conclusion

Combining automated tracking, social networks, and spatial mapping of the nest, we provide a low-dimensional representation of the multimodal interaction network of an entire honey bee colony. While many internal and external factors drive an individuals’ behavior, network age represents an accurate way to measure the resulting behavior of all individuals in a colony non-invasively over extended periods.

We use annotated location labels to select which information to extract from the social network, but stress that network age can only contain information inherent in the social network. Therefore, the predictive power of network age demonstrates that the social interaction network by itself comprehensively captures an individual’s behavior. We show that network age does not only separate bees into task groups, such as foragers and nurses, but also allows us to follow maturing individuals as they develop. A recent work derived a social maturity index in colonies of the social ant *C. fellah* (Richardson et al. 2020), similarly highlighting a strong separation of nurses and foragers in the social network and high variability in transition timing. In contrast to the fluid changes in task allocation at an individual level, we find distinct clusters of developmental trajectories at the colony level, with some groups entering critical developmental transitions earlier in life than others. Investigating the precise combination of internal and external factors that drive those transitions is a promising direction for future research.

These transition points are also reflected in changes in nest location, because spatial preferences, task allocation, and interactions are inherently coupled in honey bees. However, we show that network age is more than just a representation of location: Bees with the same network age do not necessarily share a location in the nest, and the time spent in task-associated locations is less predictive of an individual’s current and future behavior than network age. Additionally, we calculate a variant of network age that is not guided by auxiliary spatial information but instead extracts the information with the highest variance from the social networks (Network age PCA). The PCA variant is still predictive of task allocation, suggesting that location is the predominant signal in the social network. However, the higher predictive power of the CCA network age variant and the targeted embeddings indicates that there is more information in the social network that our method can extract.

In this study, we extract network age from daily aggregated interaction networks, and thereby disregard potentially relevant intraday information, which could reveal further differences between individuals (e.g., the temporal aspects of intraday interaction networks can disentangle the contribution of different modes of interactions (Hasenjager et al. 2020)). While we observe thousands of individuals and many overlapping cohorts, there is no straight-forward extension of the method to extract a common embedding of social networks that do not share individuals (e.g., over different experimental treatments or repetitions). Analyzing how network age changes within a day, over other datasets with possibly other types of interactions, or how network age shifts in response to disease pressure or experimental manipulation of age demography would be potentially fruitful areas for future investigation, as recent work has shown that there is an effect of viral infection on interaction behaviors (Geffre et al. 2020).

Network age can be repurposed and extended for other research questions: We show that (1) variants of network age capture different aspects from the social networks related to mortality, velocity, or circadian rhythms, and (2) with a subsample of only 5% of the bees in the colony, we can extract a good representation of the social network. This makes the method applicable to systems with far more individuals or with much less required experimental effort for a comparable number of individuals. Network age could be calculated in real-time, opening up a wide range of possibilities for future research: For example, it would be possible to selectively remove bees that have just begun a developmental change to determine their influence on colony-wide task allocation. Sequencing individual bees could determine how known genetic drivers of behavioral transition, like the double-repressor co-regulation of vitellogenin and juvenile hormone, are reflected in the social network. Our perspective captures both internal and external influences that impact social interactions and is thus applicable to all complex systems with observable multimodal interaction networks. Network age can be adapted to questions that explore social interaction patterns independent of age and division of labor, making it broadly applicable to any social system. As such, our method will permit future research to analyze how complex social animal groups use and modify interaction patterns to adapt and react to biotic and abiotic pressures.

## 7 Acknowledgements

We are indebted to numerous students, in particular Fernando Wario Vázquez, Franziska Lojewski, Andreas Jörg, Leon Sixt, Hauke Mönck, Maria Sparenberg, Sascha Witte, Alexa Schlegel, Mathis Hocke and Andreas Berg for providing hardware and software parts of the BeesBook system. We thank Peter Knoll and Randolf Menzel for providing the honey bee colony and valuable advice. Giovanni Galizia and Jake Graving gave critical feedback which helped improve the manuscript. Our work was supported by the HPC-Service of ZEDAT (Freie Universität Berlin) and the North-German Supercomputing Alliance who generously provided computing resources and storage capacity.

## 9 Author contributions

Conceptualization: BW, DMD, AZ, TL, MLS, KST; Methodology: BW, DMD, AZ, TL; Software: BW, DMD; Resources, supervision: TL, DB, IDC; Project administration: TL; Data curation: BW, DMD, MLS; Writing: BW, DMD, TL, MLS, KST, IDC; Visualization: BW, DMD.

## 11 Data and materials availability

The datasets generated during and/or analysed during the current study are available from the corresponding authors on reasonable request.

## Supplemental Materials and Methods

### SI 1 What is *network age*?

#### SI 1.1 Recording setup, data extraction, preprocessing

We set up our observation hive on 2016-06-24, with a queen and approximately 2,000 young bees (*Apis mellifera*) sourced from a local host colony. To obtain newly emerged bees, we incubated brood from the host colony, and later from the observation colony in an incubator at 34°C. Freshly emerged bees were marked every weekday. All bees were removed from the brood comb each day before marking, so the maximum age in each batch of bees was 24 hours. The number of bees marked per batch varied, but never exceeded 156 bees per marking. Marked bees were introduced to the colony through a backdoor entrance. After the initial marking period (27 days), the video recording was started 2016-07-24 and stopped 2016-09-19. Marking newborn bees continued approximately twice a week. A total of 3,166 individuals were marked. Bees had free access to the outside environment via a tube connected to the observation hive. The recording setup (scaffold, cameras, lighting, storage) was described previously (Wario, Wild, Couvillon, et al. 2015), however, for the experiments described here, we used custom-built IR flash circuit boards triggered by an Arduino controller, synchronized with the high-res cameras (Mönck 2016). Combs were imaged at 3 Hz, alternating between sides of the observation hive, to avoid low contrast due to backlighting.

**Figure SI 1:**
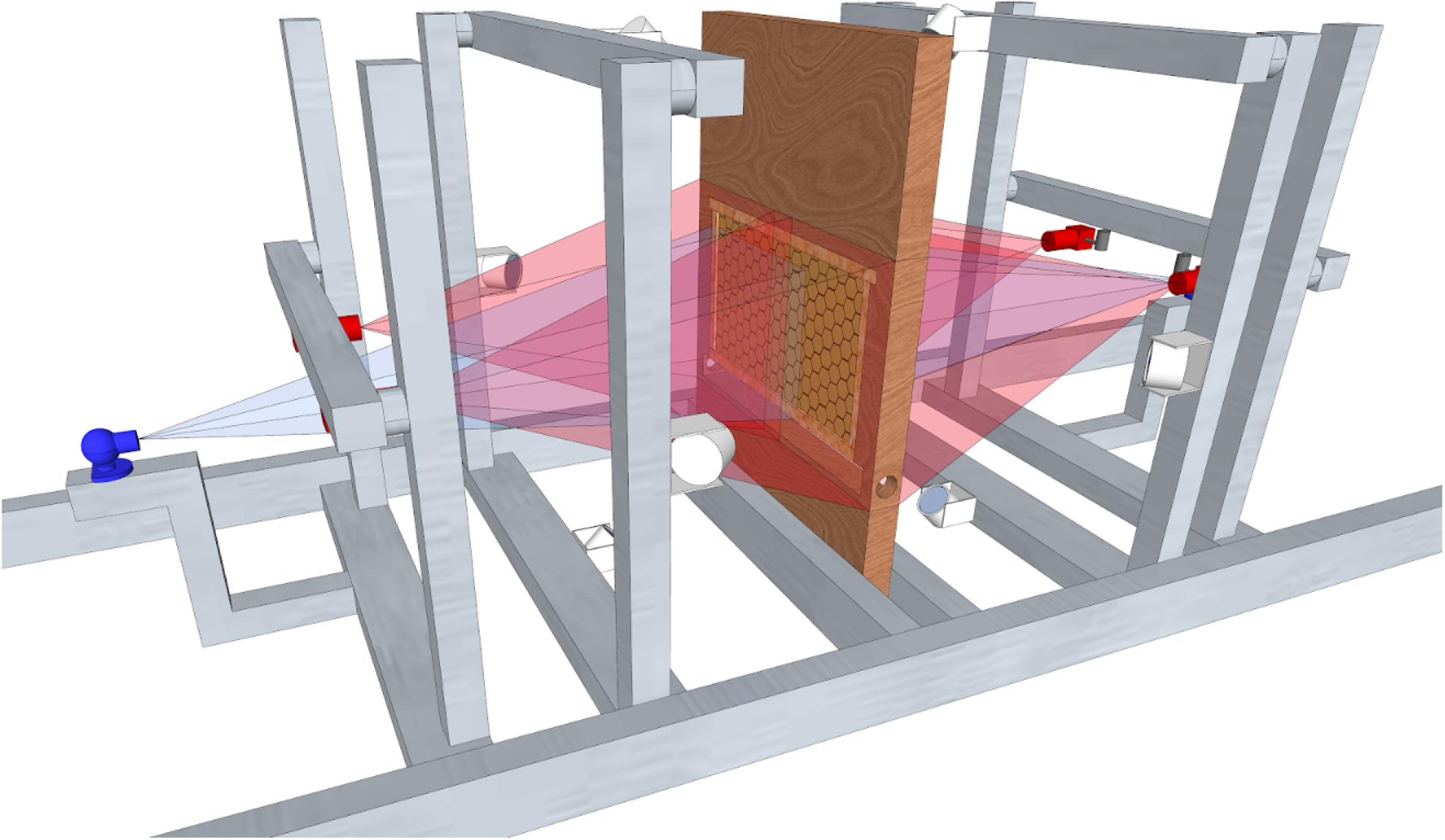
Recording setup. A one-frame observation hive stands in the center of the rig, two high-res cameras (red), and one low-res camera (blue) observe each side of the hive. The low-res cameras only observe the dance floor. Four infrared flashlights were used per side (synchronized to the high-res cameras), constant red lighting illuminated the dance floor.

A total of 46 TB of video data was recorded and continuously moved to a network storage unit at the North-German Supercomputing Alliance (HLRN). After the recording season, the data was processed to detect and decode the bee markers (Wild et al. 2018) and track these detections through time (Boenisch et al. 2018). The output data consists of timestamps, planar positions, three-dimensional rotations, IDs, and confidence scores, for the decoded IDs.

**Figure SI 2:**
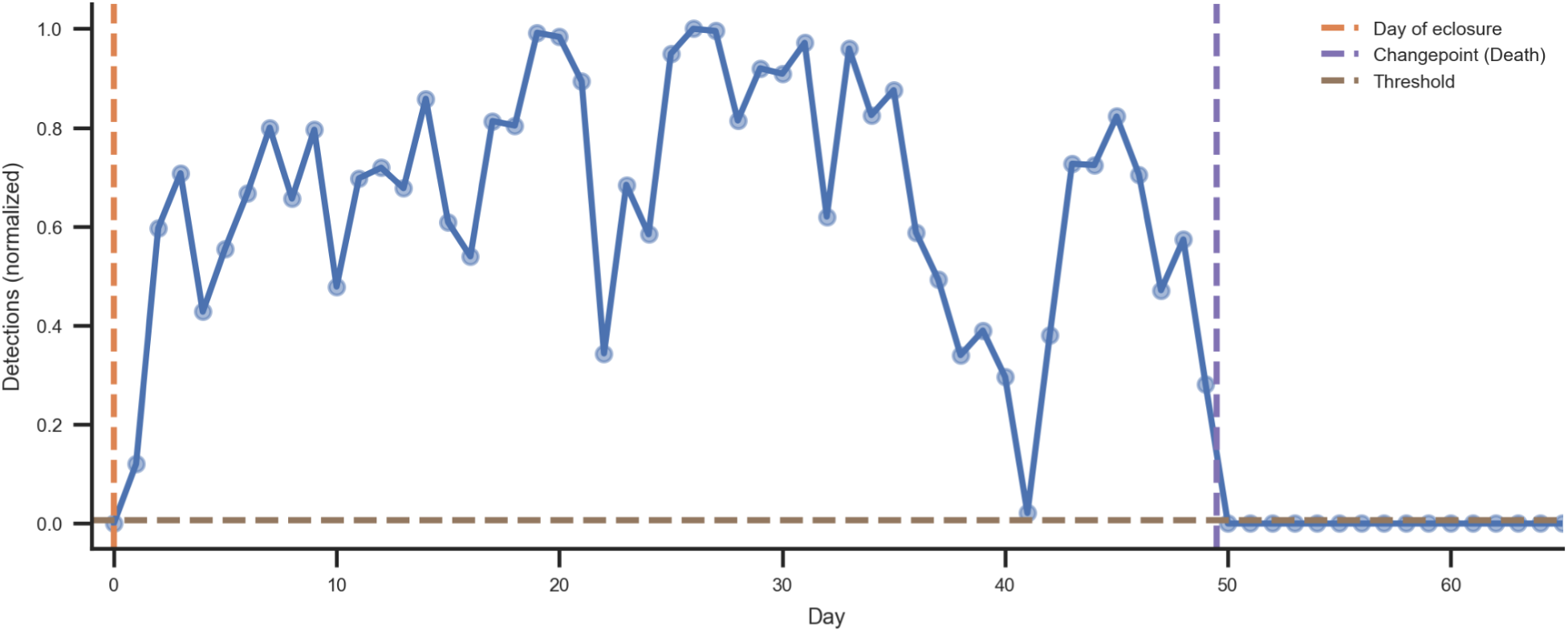
The normalized detection counts of one bee over the recording period following her introduction to the observation hive. The time of her death (purple dotted line on the right) is calculated using a Bayesian changepoint model based on optimizing a detection threshold that discriminates between days that the bee was alive and days where she was probably dead.

Two cameras were used for each side of the nest with both viewports overlapping partially. Six reference points were marked on each comb side such that four points were visible in each of the cameras’ recordings with the center two points visible in both cameras. Reference points were identified and their image coordinates were extracted manually. The coordinates were then used to calculate the homography between the comb and the image plane. The homography was then used to rectify the tracking data which translated image coordinates to a metric reference frame, i.e. the nest surface. See fig. SI 1 for a photo of the setup.

Resulting tracking data was post-processed before entering the analysis. We discarded detections with low decoding confidence, i.e. detections that the machine vision pipeline could not reliably decode. Remaining implausible detections (e.g. of IDs that had not been tagged yet) were removed in an additional filter step. The distribution of the number of detections of all IDs is strongly bimodal, the larger mode representing those bees which actually are in the observation hive, and the smaller mode representing erroneous decodings of tags which are not present on the given day. We use Otsu’s method (Otsu 1979) to automatically determine the threshold which best separates those two modes and filter out all potentially incorrect IDs. The tracking data, therefore, contains gaps due to falsely filtering out correct detections, but also due to occlusions, such as when bees inspect cells or depart the nest on foraging trips.

### SI 1.2 Bayesian lifetime prediction – Changepoint model

The death date of an individual could ideally be computed as the first date she was not detected in the hive. Unfortunately, this does not work in practice for two reasons. First, tags are sometimes incorrectly decoded, and because of the number of detections we have for each day, this means that most IDs will be detected at least a couple of times per day. Second, some bees were not visible at all on some days, even though they are not dead yet (see fig. SI 2 for an example).

We, therefore, use a Bayesian changepoint model to robustly estimate the death dates of all individuals. An individual is defined to be alive on all days since she emerged and was introduced into the colony (day *e*) up to the change point *d* = *e* + *l*, where *l* is the number of days she was alive. We use a weakly informative prior *N*(35, 50) for the number of alive days *l*. We model the probability that a bee is detected at least as often as a threshold *t* while she is alive and less often than *t* when she is not alive, using a Bernoulli distribution. We use a *Beta*(5, 1) prior for this probability because we know that, typically, an alive bee will have many detections. For the threshold *t* we use an informative *Beta*(25, 1) prior because we know that a dead bee will have very few detections, if any. Note that we normalize the detection counts to [0, 1] when fitting the model, i.e. for each bee we divide the counts of daily detections by the maximum count of detections of that tag over the entire recording period. We sample this model using pymc3 and the NUTS sampler (Salvatier et al. 2016). For each bee, we compute 2,000 tuning samples and 1,000 samples. The date of death is determined using the mean of those last 1,000 Monte Carlo samples.

### SI 1.3 Social networks

#### Proximity interaction network

Two bees were defined to be in proximity if their tags were less than 2 cm from each other (∼1.4 body lengths) over at least 0.9 seconds (three frames with our recording frame rate). We construct affinity networks based on the counts of these proximity interactions without taking the duration of each contact into account to reduce the effect of bees resting next to each other.

#### Euclidean proximity networks

Euclidean proximities were determined for each pair of bees when both bees were visible. The daily average distance *d* between two individuals was then transformed to two affinity matrices, the first derived by applying a Gaussian similarity function *d*′ = (*d*^2^*/*2*γ*^2^) with *γ* = *max*(*D*)*/*4 and the second by subtracting from the maximum distance (*d*′ = *max*(*D*) − *d*). *D* is the matrix of all daily average distances on the same day.

#### Trophallaxis networks

We constructed an interaction network representing trophallaxis interactions (food exchange). To filter our data to detect trophallaxis events, we use a two-step approach. We first use a fast logistic regression with low precision to discard most of the non-trophallaxis encounters. We then use a slower convolutional neural network to further refine the results with higher recall.

To train the two classification models, we manually labeled bee interactions in our dataset by observing video sequences. To increase the fraction of positive events, we queried our data for bees that are close to each other, and approximately facing each other.

This ground truth data contains 140 trophallaxis events out of the distinct 2,651 events in total. For some events, we annotated a begin and end timestamp and could, therefore, use multiple frames for the training. In total, we had 25,835 training samples, each consisting of a pair of bee IDs, a timestamp, and a label (trophallaxis / not trophallaxis).

Because the prefiltering of the training data can introduce a sampling bias we created another test set by labeling all possible interactions in 33 randomly sampled frames, containing a total of 15 trophallaxis events and 39,051 negative events (we use every pair of bees with a thorax distance of less or equal than 3 cm as a possible candidate). This test set represents our data distribution without any bias.

In the classification step, we look at all pairs of bees with their thoraxes at a distance between 0.731 cm and 1.204 cm (i.e. the 99th percentile of the positive events in the training data) together in a frame. For a pair of bees (*i, j*) we have the locations of the thorax on the hive in millimeters (*xy*_*i*_, *xy*_*j*_) and their orientations (*α*_*i*_, *α*_*j*_). We calculate the approximate head position *h*_*i*_ as *xy*_*i*_ + *d* [*cos*(*α*_*i*_), *sin*(*α*_*i*_)] where *d*=3.19 mm. We calculate their relative orientation as [*cos*(*α*_*i*_), *sin*(*α*_*i*_)] [*cos*(*α*_*j*_), *sin*(*α*_*j*_)]^*T*^. We then perform logistic regression on the euclidean distance of their thorax locations, the euclidean distance of their head locations, and their relative orientation.

The logistic regression was trained on the manually labeled samples, setting the threshold to get a recall of 85% at a precision of 21% (on a 20% validation test). This regression discards 62% of the true negative samples (i.e. the specificity). For the remaining data points, those that were classified as possible trophallaxis by the first classifier, we extract trajectories of both bees for around 5 seconds (15 frames) around the possible trophallaxis events. We then use a convolutional neural network, again trained on the manually labeled data.

Evaluated on the test set, the two combined filters yield a recall of 60% at a precision of 47%, discarding 99.97% of negative samples (i.e. the specificity).

#### Interaction effect networks

For each proximity interaction with a duration no longer than 60 seconds (to exclude bees resting next to each other) and with a minimum gap of at least 5 seconds since the last interaction of the same two individuals, we compute the difference in mean velocities within 30 second time windows before and after the interaction. This is done for both partners, and so we derive four networks based on the mean and cumulative changes, each split into negative and positive values. We use separate networks for the positive and negative values because this allows us to define affinity matrices which can only have positive edge weights.

#### Temporal aggregation and post-processing of networks

Time-aggregated networks were constructed by defining the weighted edge strength as the number of times two individuals were in proximity or engaged in trophallaxis. The networks were aggregated over 24 hours without overlap. Edges in both networks are undirected. For subsequent analyses, all networks were represented as a square adjacency matrix with each element *i, j* representing the affinity of bee *i* with bee *j* on this day, given by the interaction mode (e.g. for the network of trophallaxis counts, a high value represents many trophallaxis interactions between the two individuals). Each matrix is then preprocessed using a rank transform and normalized such that 0 represents the lowest affinity and 1 the highest affinity. Ties are resolved by assigning the same rank to identical affinities.

### SI 1.4 Nest area mapping and task descriptor

We manually outlined the capped brood area and visible honey storage cells for every day in background images from 2016-07-30 until 2016-09-05. To obtain the open brood area, we calculated the area of the comb that would become capped within 8 days. We extracted the background images by extracting the first frame from every video we recorded over a specific day (approximately one image every 5.6 minutes), and then applying a rolling median filter with window size 10 to these images. We then calculated the modal pixel value, for every pixel, over all the median images. For each side of the comb, we stitched together the background images from the two cameras on that side.

To get the approximate location of the dance floor, we used the detected waggle runs of our waggle dance detection system (Wario, Wild, Rojas, et al. 2017) that had high confidence (>=0.9) (see fig. SI 3). As nearly all waggle detections happened on one side of the comb, we exclusively labelled this area as the dance floor. We then fitted an ellipse to the detections using scikit-image (van der Walt et al. 2014), which we scaled manually to not intersect with the exit area. The dance floor area was consistent throughout the experiment, so in cases where we did not have waggle dance data for a given day, we interpolated the dance floor area over the adjacent days. Finally, we used a kaiser window applied over the consecutive days (window size=5, beta=5) to smooth the annotations. We considered the region 7.5 cm around the exit tube as the nest region close to the exit.

To generate a task descriptor for every bee, we fetched one high confidence detection (>0.9) per bee for every minute of a day. We then counted how many of these detections per bee fell into the annotated regions. Then we normalized these counts per bee to 1 by dividing through the sum. This descriptor, therefore, contains the fraction of time each individual spends in each of the annotated regions. Data points outside of the annotated areas are ignored for this descriptor but are used in all other parts of this work. For all evaluations, we consider the brood area region to be the sum of the annotated open and closed brood cell regions. See Fig S004B for an example of the annotations.

**Figure SI 3:**
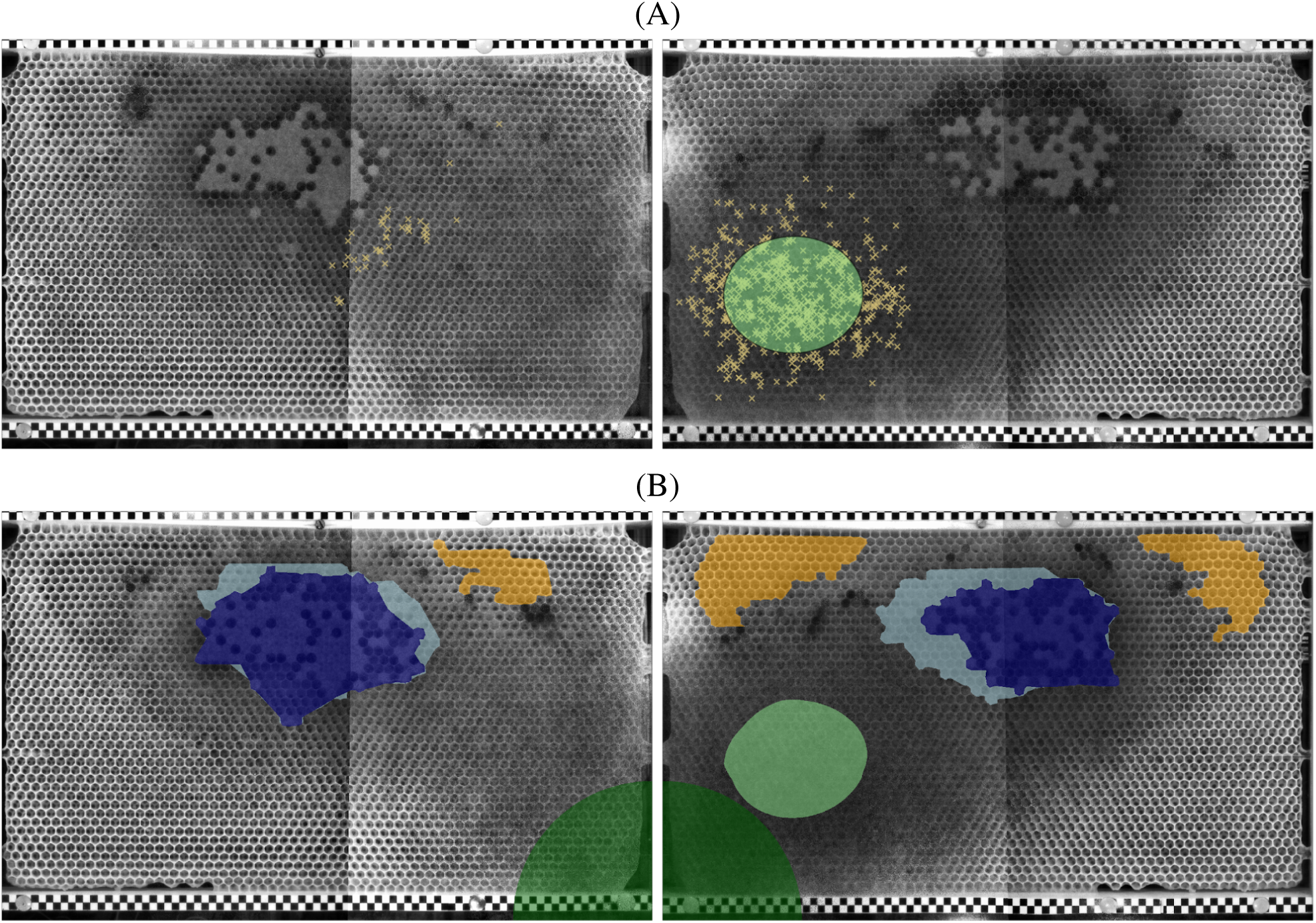
Ground truth data for the location-based task descriptors. (A) Background-subtracted images of the two sides of the observation hive. The images from the two cameras per side have been stitched together. The markers show the location of the high-confidence waggle run detections for 2016-08-18. Nearly all are on one side of the comb. The green region indicates the dance floor location. (A) Annotations for 2016-08-18, used to generate task descriptors for every bee (dark blue: capped brood, light blue: open brood, yellow: honey, light green: dance floor, dark green: exit region).

### SI 1.5 Network age method – From networks to spectral embeddings to CCA

Network age is derived from the raw interaction matrices using spectral decomposition and canonical correlation analysis (CCA). For each day and interaction mode, the graph of interactions between all bees that were alive (see section SI 1.2 for the definition of alive bees) on that day is retrieved as an adjacency matrix as described in section SI 1.3.

For each preprocessed affinity matrix, spectral embeddings (Belkin et al. 2003) are calculated using the python package scikit-network. We compute the first eight embedding dimensions (see section SI 2.2 for an evaluation of the performance of different numbers of embeddings).

For the non-symmetric interaction effect matrices, we use bispectral decomposition (Abdi 2007) to obtain one set of embeddings each for the rows and for the columns, to represent the two directions of an interaction.

For different days the eigenvectors of the embeddings and therefore the embedding values themselves can have an inverted sign. To correct this, we flip the sign of the values if the Spearman correlation between consecutive days is negative.

For every day, we now have a high-dimensional embedding per bee. We reduce the dimensionality further by applying canonical-correlation analysis (CCA). We use CCA to find a linear transformation of the network embeddings to a three-dimensional vector that maximizes the correlation to a projection of the bees’ task descriptors (as introduced in section SI 1.4). We use the CCA implementation in scikit-learn (Pedregosa et al. 2011). We use the first dimension of this vector as *network age* throughout this paper, but also evaluate multidimensional variants (Network age 2D, Network age 3D) in sections SI 2.4, SI 2.6 and SI 4.3.

For every dimension and day, we use robust scaling based on the 5th and 95th percentile of the network age distribution. A network age of zero corresponds to the 5th percentile and 40 corresponds to the 95th percentile. This stabilizes the distribution over time and also maps the values to a range comparable with the biological age of honey bees. We note that this scaling slightly improves the prediction of task allocation, but that the method also works without it. We enforce that the 5th percentile of network age corresponds to bees with a lower biological age than the 95th percentile such that biological age and network age have the same directionality.

### SI 1.6 Network age method – Unsupervised variant using Principal Component Analysis

We also calculate a variation of network age that does not require the annotated location descriptors. Instead of applying CCA to the concatenated spectral embeddings (see section SI 1.5), we instead use Principal Component Analysis (PCA) to reduce the dimensionality. This unsupervised network age still predicts task allocation better than biological age (see sections SI 2.1 and SI 2.4 for details).

### SI 1.7 Network age distributions over time

While the network age distribution stays within similar bounds over time, we find that the shape of the distribution (e.g. the proportion of individuals in the lower vs higher percentiles) changes significantly over the experimental window, likely reflecting changes in task allocation (e.g. more individual engaged in brood care initially and more foragers after most brood cells are capped). See fig. SI 4 for distributions over time.

We find no strong evidence of correlations of these changes in distribution with external factors such as the weather, but this may be due to the limited size of the recording window and is an interesting direction for potential future research.

### SI 1.8 Location heatmaps

To visually assess how distinctive biological age and network age are with respect to the spatial distribution of individuals, we computed location heatmaps for single-age cohorts tracked over the experimental window (25 days). For each individual within the cohort, we collected their daily biological age, network age, and their positions on the comb (as in section SI 1.4). Positional data for each day (a *bee-day*) was then assigned to a heatmap as follows: For the age heatmaps, we manually defined age intervals based on the age-thresholds given by Seeley (Seeley 1982), with an additional split for middle-aged bees for which we observed a high variance. We then assigned bee-days according to these thresholds based on the bees’ ages on the different days. The network age thresholds for the plots have been set in a way to keep the number of bee-days the same for the two plots in each column and can, therefore, differ for different cohorts. For example, the cohort introduced on 2016-08-01 consisted of 123 bees. In the heatmap for age 0-11, there would be N=1394 bee-days over the focal period in which the bees were below 12 days old (some bees having died). Then, we calculated the first network age thresholds in such a way, that they would also include 1394 bee-days.

**Figure SI 4:**
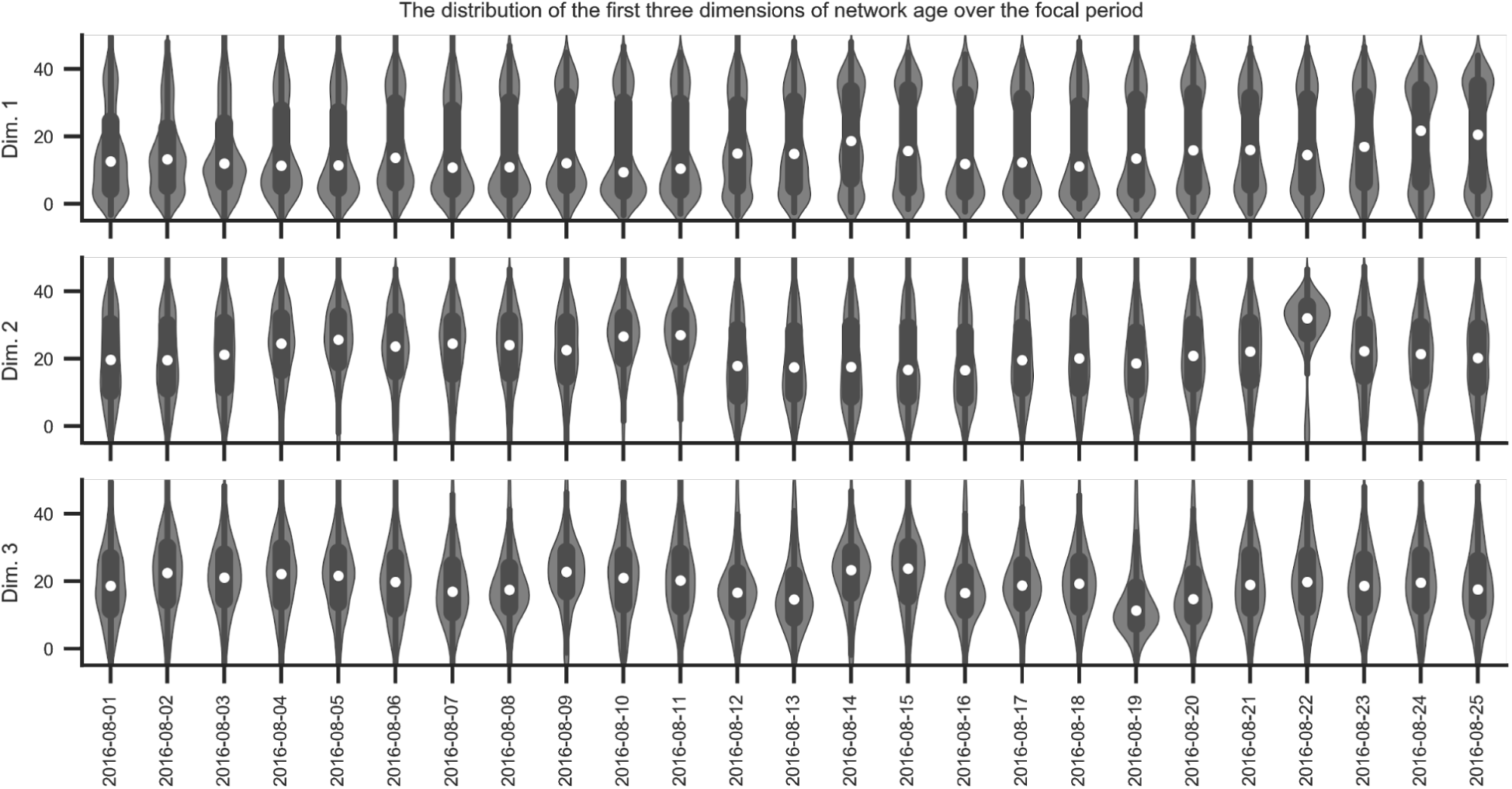
Violin plots of network age distributions over the focal period. The first three dimensions of network age are depicted. Each violin comprises one dimension of the network age values of all individuals on a given day (boxes: center dot, median; box limits, upper and lower quartiles; whiskers, 1.5x interquartile range).

The shaded areas depict density percentiles (brightest to darkest: 99%, 97.5%, 95%, 80%, 70%, 20%).

See figs. SI 5A and SI 5B for heatmaps showing the distributions of different example cohorts that we introduced at the beginning of the focal period and fig. SI 5C for all bees.

### SI 2 Network age correctly identifies task allocation

#### SI 2.1 Variability of task prediction accuracy over time

The task allocation prediction scores are not constant over time for biological age, network age, and the unsupervised (PCA) variant of network age. Here, we show McFadden’s Pseudo *R*^2^ scores for individual days. While there is variance in the predictability using network age, we note that on all dates, the models based on network age yield a higher score than the models using biological age. On all days, the unsupervised PCA variant is more predictive than biological age with comparably small differences to the CCA variant. See fig. SI 6 for the distribution and timeline.

### SI 2.2 Effect of subsampling bees, the dimensionality of spectral embeddings, and only using proximity information on task prediction accuracy

Network age can be computed with a much smaller proportion of marked bees, hence allowing one to study much larger colonies, or reducing the effort for marking individuals. To quantify the effect of sparse sampling, we perform the following analysis. We subsample the individuals entering the analysis and compute interaction networks only for those selected individuals, i.e. only interactions between individuals of this subset are recorded. We then compute spectral factors and network age as described in section SI 1.5. We find that network age is remarkably robust in this setting. With 5% of the bees tagged, the proportion in variance explained using network age is comparable to the case of using all individuals. Even with only 1% of the individuals used in the analysis, network age still is a better predictor of task allocation than biological age. This suggests that a much larger colony could be analyzed with this method without additional effort, as long as a representative subset of the bees is tracked. See fig. SI 7A for the influence of different subsampling fractions on the results.

**Figure SI 5:**
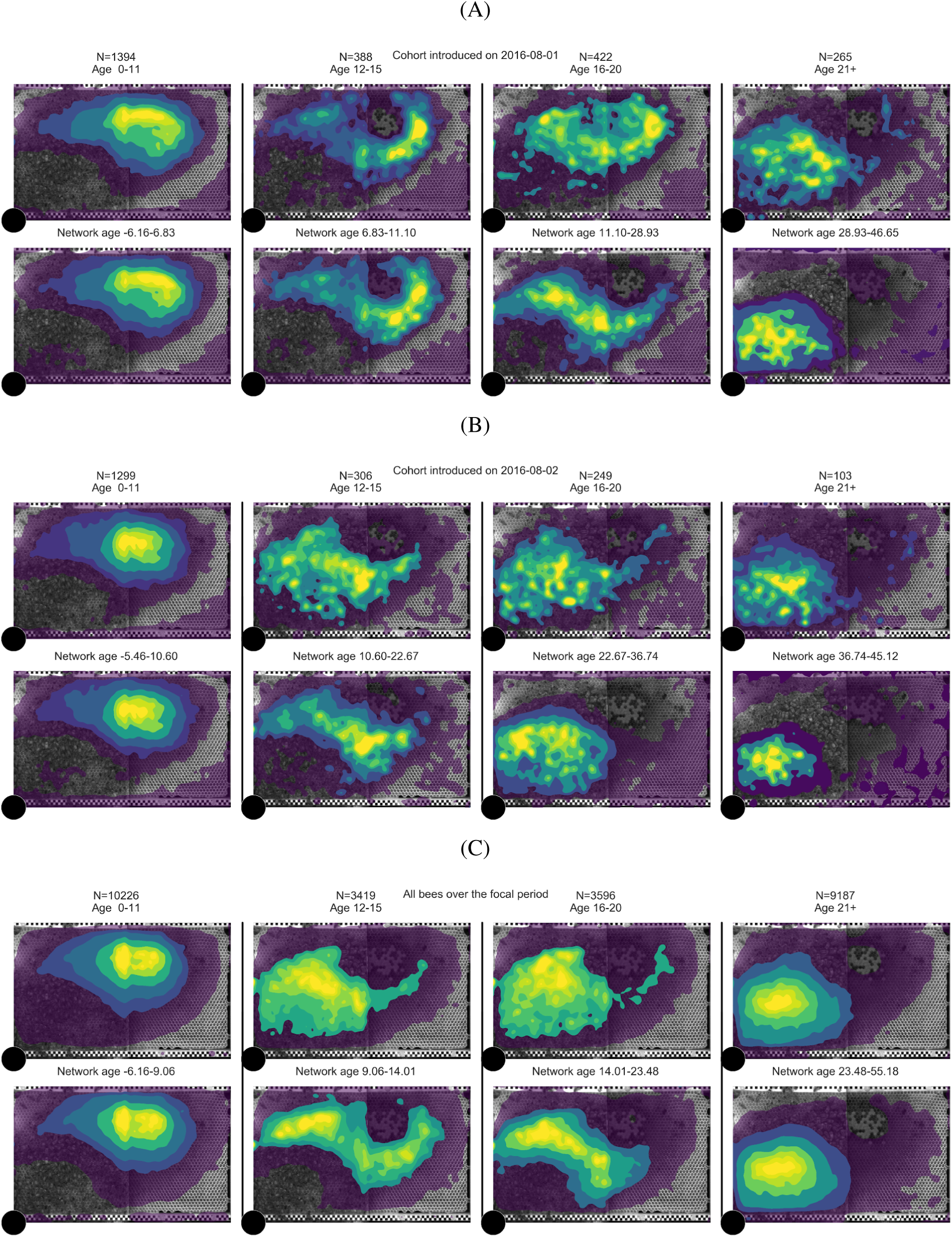
Spatial heatmaps of bees grouped by biological age and network age. (A) Cohort of 2016-08-01 (N=123 bees) (B) Cohort from 2016-08-02 (N=119 bees) (C) All bees over the focal period (N=1912 bees)

**Figure SI 6:**
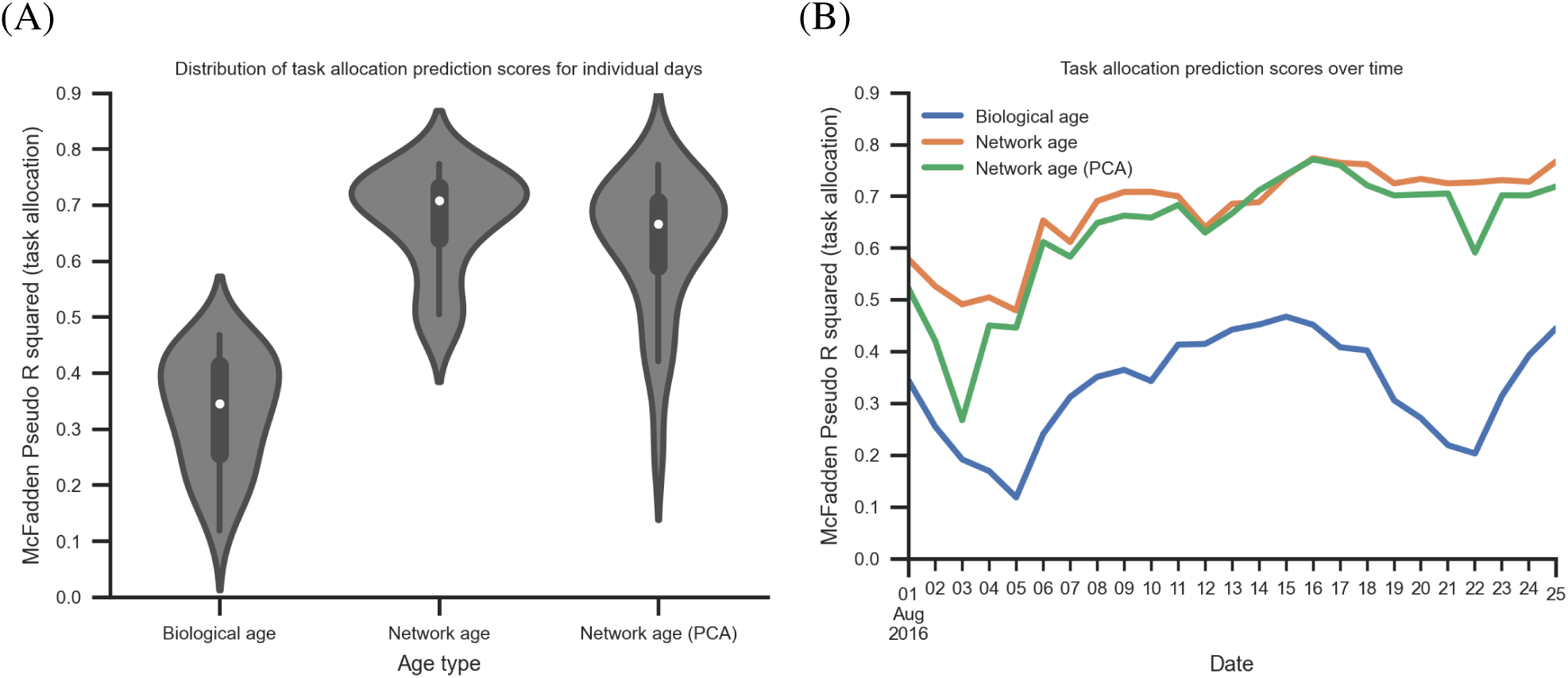
Stability of task allocation prediction over time. (A) The distribution of McFadden’s *R*^2^ for task allocation prediction for the different days using either network age, the unsupervised network age (PCA), or biological age as predictors (boxes: center dot, median; box limits, upper and lower quartiles; whiskers, 1.5x interquartile range). (B) The same data points plotted over time.

**Figure SI 7:**
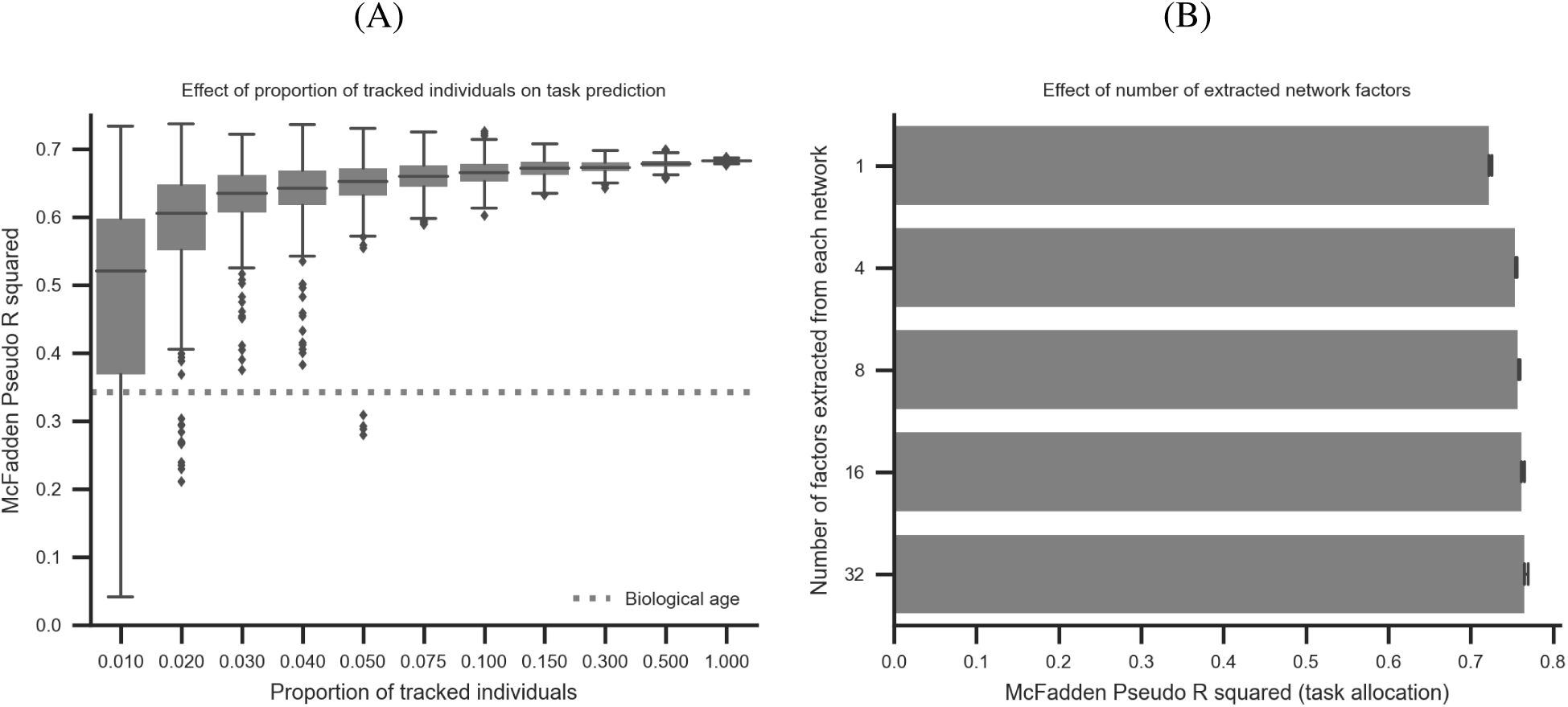
Effects of subsampling the individuals and the number of spectral factors from each interaction network on the task allocation prediction. (A) McFadden’s *R*^2^ for different fractions of subsampled individuals. While having all bees individually marked and tracked yields the highest median score, tracking only 5% of the individuals yields comparable results when drawing a representative subset. Each box comprises 128 bootstrap runs (center line, median; box limits, upper and lower quartiles; whiskers, 1.5x interquartile range; points, outliers). (B) The influence of the number of dimensions per interaction network extracted using Spectral embedding before applying the CCA on the task allocation prediction scores (using network age 3D). While more dimensions enable the CCA to extract more relevant information, the overall improvement is small. The bars give the mean of 128 bootstrap samples per dimensionality and a bootstrap sampled 95% CI of the mean.

Likewise, we also find that the number of spectral factors per interaction mode matrix does not have a strong effect on the proportion of variance explained by the network age extracted from those factors. Using more factors is computationally more expensive and produces statistically better embeddings. However, network age computed using 32 factors explains task allocation only marginally better compared to network age computed using 4 factors (see fig. SI 7A).

We also evaluate variants of network age that can be derived using only the spatial information of detected individuals. To that end, we exactly follow the procedure outlined in section SI 1.5, but discard all interaction types except for either spatial proximities (*Euclidean proximity* variant, see *Euclidean proximity networks* in section SI 1.3) or interaction events derived from spatial proximity (*Proximity events* variant, see *Proximity interaction network* in section SI 1.3). We find that network age derived from all interaction types outperforms these proximity based variants in terms of task allocation, but the variant based on proximity events is only marginally worse (see section SI 2.4) and could be used in future studies if only proximity data is available.

### SI 2.3 Spatial separation of bees with similar network age

While the network age of bees is correlated with their locations, we do find examples of bees that have approximately the same network age but occupy different locations on the comb (see fig. SI 8 for an example). We also see that, while two bees having a similar network age is correlated with an increase in proximity interactions, there are many bees with close network ages that do not encounter each other inside the hive (see fig. SI 9).

**Figure SI 8:**
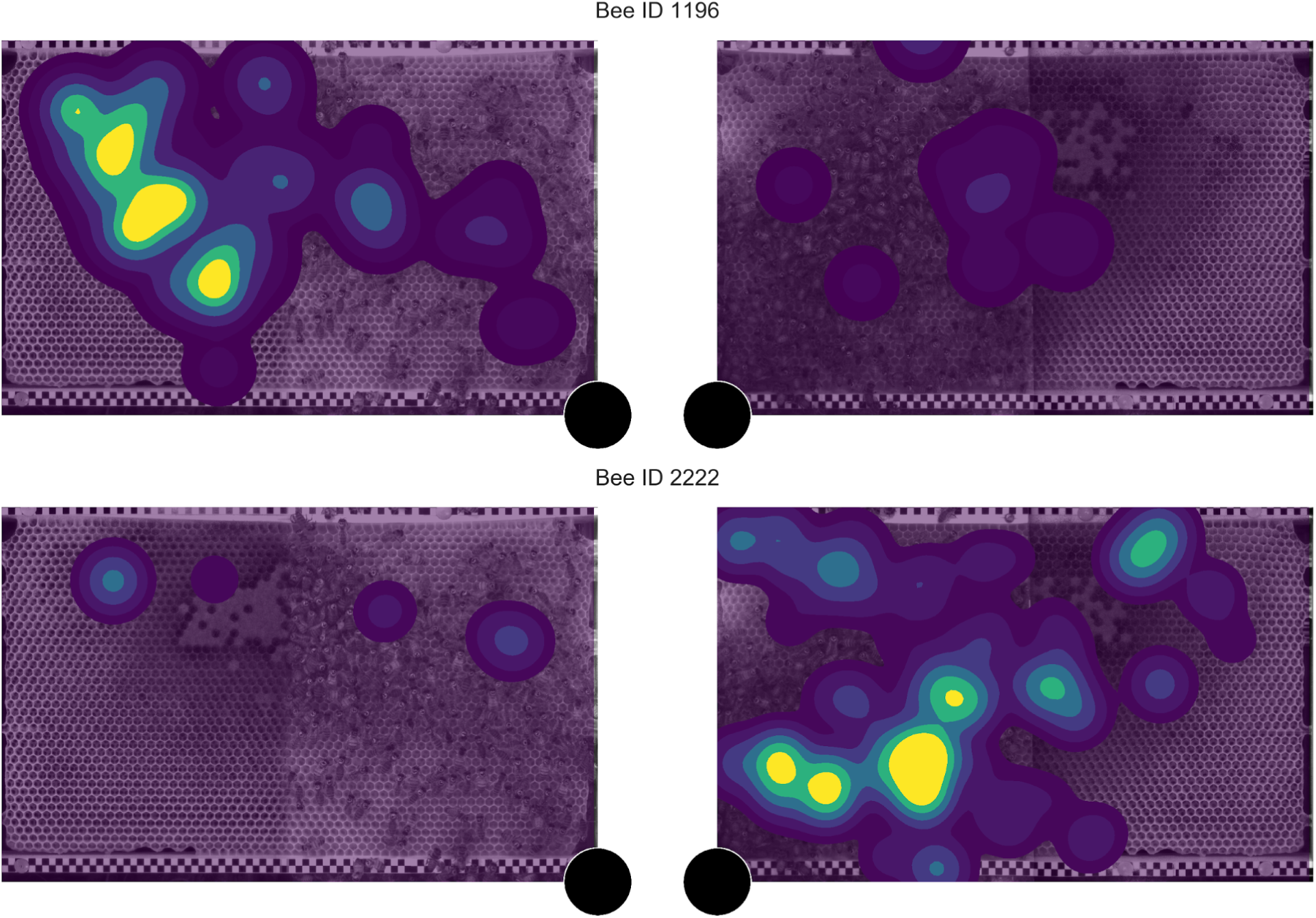
Two bees on 2016-08-05 with similar network ages despite occupying different sides of the comb. The bee with the ID 1196 is 22 days old and has a network age of 15.09. Her heatmap consists of 1189 samples, each corresponding to one minute on the focal day. The bee with the ID 2222 is 8 days old and has a network age of 14.34. Her heatmap consists of 1284 samples, each corresponding to one minute on the focal day. The black circle marks the location of the entrance.

**Figure SI 9:**
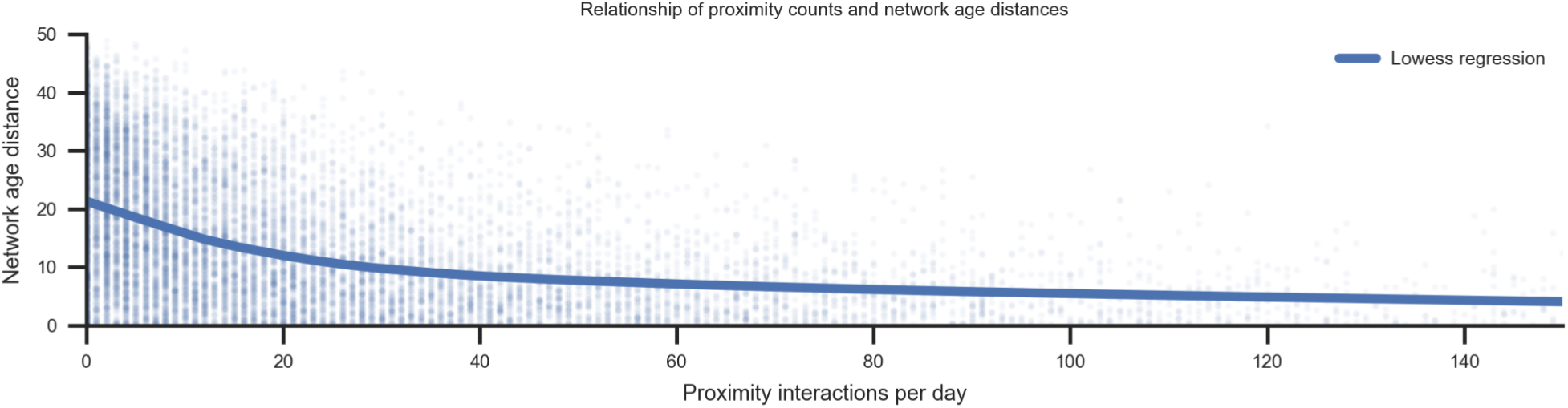
The relationship between distance in network ages and number of proximity interactions for pairs of bees. Close network ages generally mean more interactions. However, there are many individuals with a close network age and very few interactions. Each point indicates one pair of individuals on one day. The line is given by a lowess regression.

### SI 2.4 Task prediction models and bootstrapping

To evaluate how well biological age and the different variants of network age represent an individual’s task allocation, we use these measures as features to predict the proportion of time individuals spend in the brood area, dance floor, honey storage and near the exit (see section SI 1.4 for details on the nest area mapping). We evaluate the areas individually and in combined models. We evaluate different complexities of models (linear vs. nonlinear) and different independent variables (e.g. network age and biological age).

To test different complexities of the relationships, we evaluate both a generalized linear model (GLM, the default model) and a small neural network consisting of two fully connected layers (listed as *nonlinear* in table 1) for each of the combinations of independent and dependent variables. The hidden layer of the neural network has a dimensionality of 8 and uses tangens hyperbolicus as its nonlinearity.

To evaluate the performance of the models for each area separately we select a sigmoid as the link function of the GLM and the activation function of the neural network’s last layer. We then optimize and calculate the likelihood of the data assuming a binomial distribution.

We also evaluate both models to simultaneously predict all four values of an individual’s task allocation distribution. To this end, we choose a softmax function as the link function of the GLM and the neural network’s final activation function. We then optimize and calculate the likelihood of the data assuming a multinomial distribution.

For all the combinations of independent and dependent variables, we repeat the described procedure for 128 bootstrap samples. For each model, we retrieve the final likelihood of the data *L*_1_. We use PyTorch (Paszke et al. 2019) and the L-BFGS optimizer to obtain maximum likelihood estimates of the models. We also always fit a null-model only consisting of the intercepts and retrieve its likelihood *L*_0_. For each model and bootstrap iteration, we calculate McFadden’s pseudo *R*^2^ (McFadden 1973) as 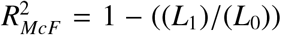. We then calculate the median and 95% confidence intervals from these bootstrap samples. See table 1 for an overview of the results for all evaluated models. We test the significance of these results separately with the tests described in section SI 2.6.

### SI 2.5 Forager groups experiment and data recording

**Figure SI 10:**
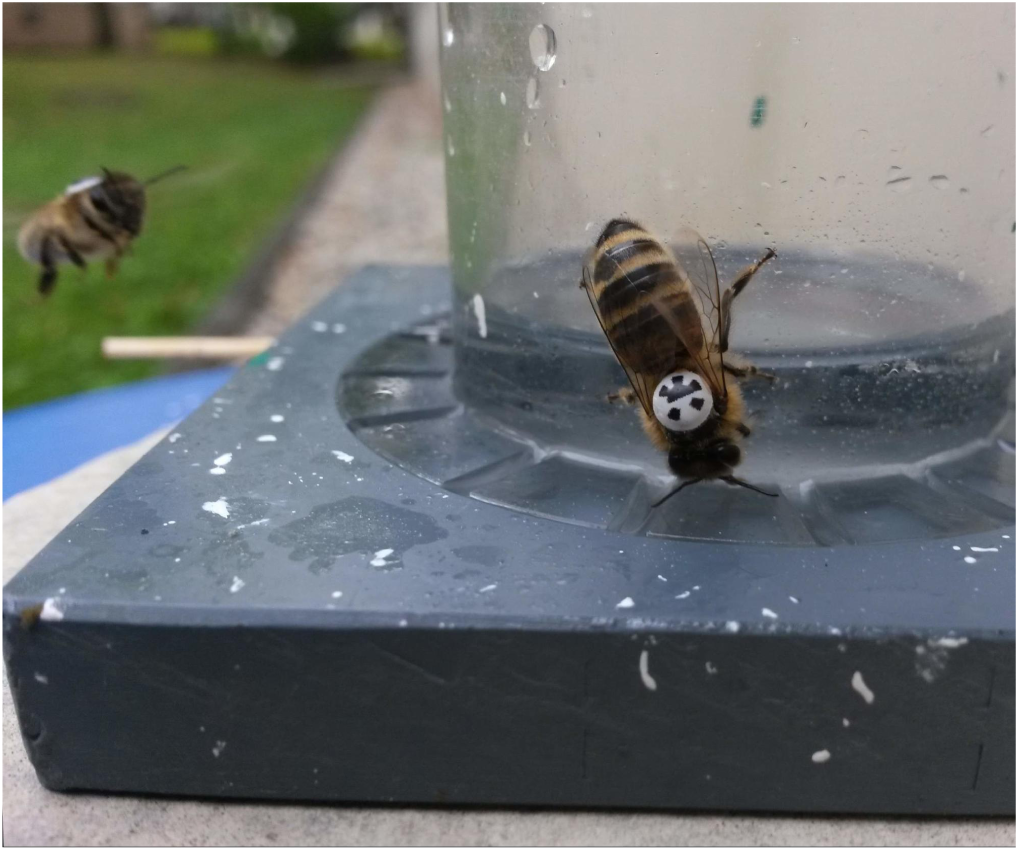
Individually marked bees at a feeding site.

**Figure SI 11:**
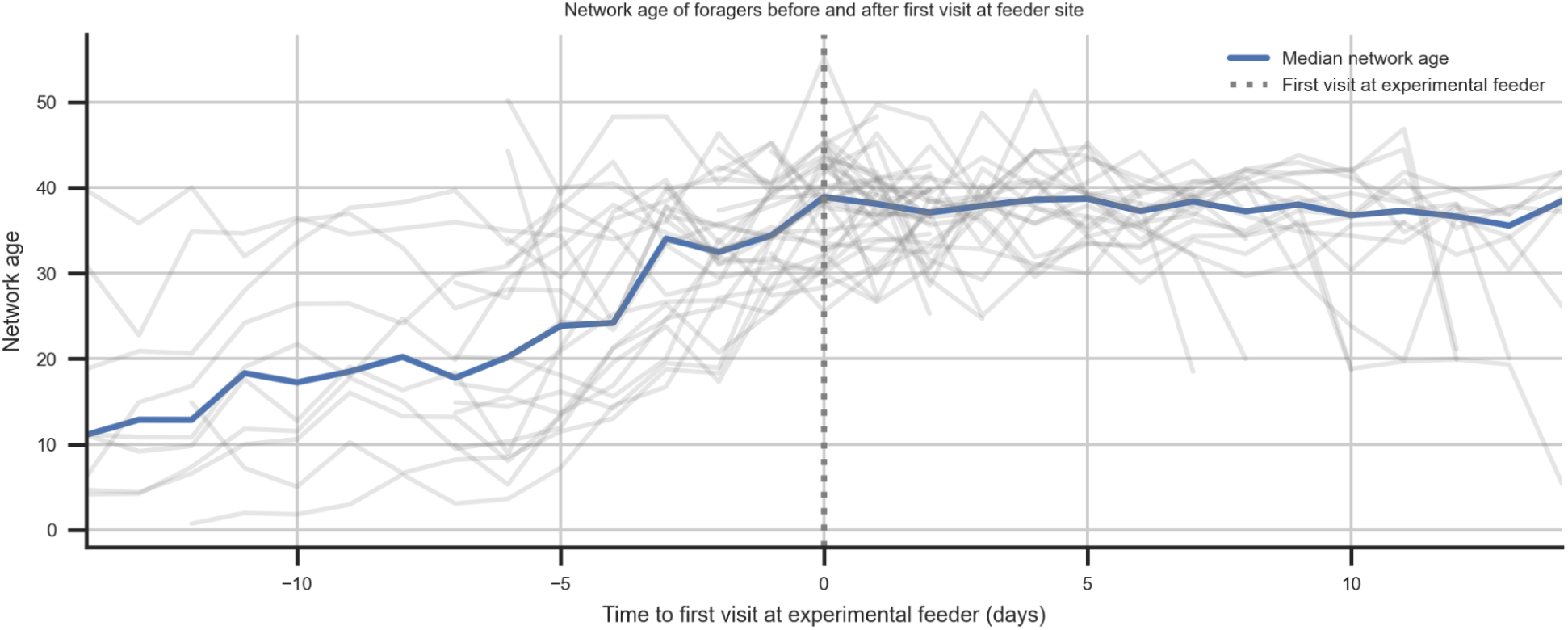
Network age development of new foragers. For every visiting bee, their first visit to a feeding station was recorded (age 12-40 days, N=40). The plot shows the network age for these bees over the previous and next days centered on their first visit. The grey lines give the network age of individual bees and the blue line indicates the median.

Between 2016-07-28 and 2016-08-22, foragers were trained to a feeder (see fig. SI 10 for a photo of the feeder) offering unscented sucrose solution by gradually moving it from the colony (52.457 032, 13.296 635) to a sequence of locations *F1* to *F4* (see table 2). For days over which the feeder was moved, high sugar concentration was used and iteratively changed to control the number of new foragers. Once the final locations were reached, the feeder offered the highest concentration for 1-2 hours per day. After a minimum of three days, training to the next location in the list was resumed. We photographed all bees landing at the location and manually transcribed the identity and time of arrival. A list of foragers visiting the feeder is given in table 3. The network age values around the day we first observed each bee at the feeding side is given in fig. SI 11.

**Table 2:** GPS coordinates and distances from the hive for the feeders used in the forager group experiment.

| Feeder | GPS coordinate | Distance from hive |
| --- | --- | --- |
| F1 | 52.455726, 13.294224 | 217 m |
| F2 | 52.454866, 13.293092 | 338 m |
| F3 | 52.454547, 13.291737 | 430 m |
| F4 | 52.453682, 13.292841 | 451 m |

**Table 3:** Tag IDs and locations observed forager bees during the forager group experiment.

| Date | Location | Forager bees observed at feeder |
| --- | --- | --- |
| 2016-08-01 | Hive-F1 | 135, 199, 217, 220, 228, 233, 253, 319, 392, 539, 727, 818, 845, 860, 1110 |
| 2016-08-02 | F1 | 135, 217, 233, 253, 648, 845, 860 |
| 2016-08-03 | F1 | 135, 228, 233, 253, 392, 648, 845, 860 |
| 2016-08-04 | F1 | 220, 228, 233, 392, 648, 710, 885 |
| 2016-08-05 | F1 | 135, 220, 228, 233, 392, 644, 648, 710, 769, 885, 931 |
| 2016-08-08 | F1-F2 | 199, 319, 337, 392, 555, 644, 710, 769, 885, 1593 |
| 2016-08-09 | F1-F2 | 199, 319, 337, 392, 555, 644, 648, 710, 769, 785, 885, 912, 1122, 1362, 1518, 1593, 2031 |
| 2016-08-10 | F1-F2 | 199, 319, 337, 392, 648, 769, 885, 1122, 1362, 1593, 2031 |
| 2016-08-11 | F2 | 199, 319, 337, 392, 644, 769, 785, 885, 1122, 1362, 1518, 1593, 2031 |
| 2016-08-12 | F2 | 199, 319, 337, 392, 769, 785, 885, 1122, 1362, 1518, 1593, 2031 |
| 2016-08-14 | F2 | 199, 319, 337, 392, 555, 644, 710, 769, 785, 885, 1232, 1362, 1518, 1593, 2031 |
| 2016-08-16 | F2 | 199, 319, 337, 710, 769, 885, 912, 1122, 1362, 1518, 1593, 2031 |
| 2016-08-17 | F2 | 199, 319, 337, 644, 710, 769, 885, 912, 1122, 1518, 1593, 2031 |
| 2016-08-18 | F2 | 199, 319, 337, 644, 769, 885, 912, 1122, 1518, 2031 |
| 2016-08-19 | F2 | 319, 710, 885, 1593, 2031 |
| 2016-08-22 | F2-F3 | 319, 644, 710, 885, 1593, 2031 |
| 2016-08-23 | F3 | 1180, 1197, 1232, 1471, 1593, 1662, 1714, 1799, 2031, 2106, 2984 |
| 2016-08-24 | F3 | 1180, 1197, 1232, 1471, 1593, 1714, 1793, 1799, 2031, 2106, 2984 |
| 2016-08-25 | F4 | 1180, 1197, 1714, 1793, 1799, 2031, 2106 |

### SI 2.6 Statistical comparison of models

We use bootstrapped confidence intervals of the effect strength to investigate whether a model based on one feature (e.g. network age) explains the dependent variables (e.g. task allocation distributions) significantly better than the same model based on a different feature (e.g. biological age). Additionally, we use a likelihood ratio *χ*^2^ test to answer whether one feature (e.g. network age) provides additional information over biological age in a combined model.

#### Bootstrapped confidence intervals of the effect strength

We draw 128 bootstrap samples of the combined daily bee data. For each sample, we calculate either the McFadden’s pseudo *R*^2^ in the case of the task allocation models (see sections SI 2.4 and SI 4.4) or the *R*^2^ in the case of the other measures (see sections SI 4.3 and SI 4.5) for both a model based on biological age and the independent variable we want to compare with (e.g. network age). For each of these paired samples, we calculate the difference in scores of the two models. From these 128 differences, we calculate a two-sided 95% confidence interval of the effect strength. If the null hypothesis (difference in scores is zero or less) is not contained in the confidence interval, we can reject the null hypothesis at an alpha level of 2.5%.

#### Likelihood ratio test

As the likelihood ratio test requires a nested model for testing, we compare a model based solely on biological age with a model based on a combination of biological age and the independent variable we want to compare with (e.g. network age).

We fit each model to the data and calculate the likelihoods of the data under the fitted models (*L*_1_ for the combined model and *L*_0_ for the model based on biological age). The likelihood ratio is given by *LR* = 2 *ln*(*L*_0_*/L*_1_). If the null hypothesis that the models are equal were true, *LR* would approximately follow a chi-squared distribution with k degrees of freedom (with k=4 in the case of the task allocation model from section SI 2.4 and k=1 in case of the general regression model for sections SI 4.3 and SI 4.5). We use the cumulative density function of the chi-squared distribution to calculate the p-value.

### SI 3 Developmental changes over the life of a bee

#### SI 3.1 Repeatability

We calculate the repeatability *R* of the network age consisting of repeated measurements over several days of an individual *I* as *R*(*I*) = *Var*_*p*_*/*(*Var*_*i*_ + *Var*_*p*_) with *Var*_*i*_ being the variance of the network age of an individual measured over the available days and *Var*_*p*_ being the variance of the mean network ages of a control group. The control group consists of all bees inside the same age span as *I* on all days on which the network age values for *I* were collected. A repeatability close to 1 means that the individual variance is low compared to the population variance. A repeatability close to 0 means that the individual variance outweighs the population variance.

### SI 3.2 Network age transition clustering

In order to cluster the transitions of different individuals in a cohort, we first collect the network ages for every individual in a feature vector where each entry corresponds to the individual’s network age for one day. We then do a linear inter- and extrapolation for missing values (e.g. due to absence or the individual dying). For the cohort of bees, we calculate the euclidean distance between each individual’s feature vector. Then we perform a hierarchical clustering using Ward’s method (Ward 1963) using the Python library scikit-learn (Salvatier et al. 2016) and extract the first three clusters. See fig. SI 12 for an example of the clustering. See fig. SI 13 for the network age development of different cohorts and fig. SI 14 for all bees.

**Figure SI 12:**
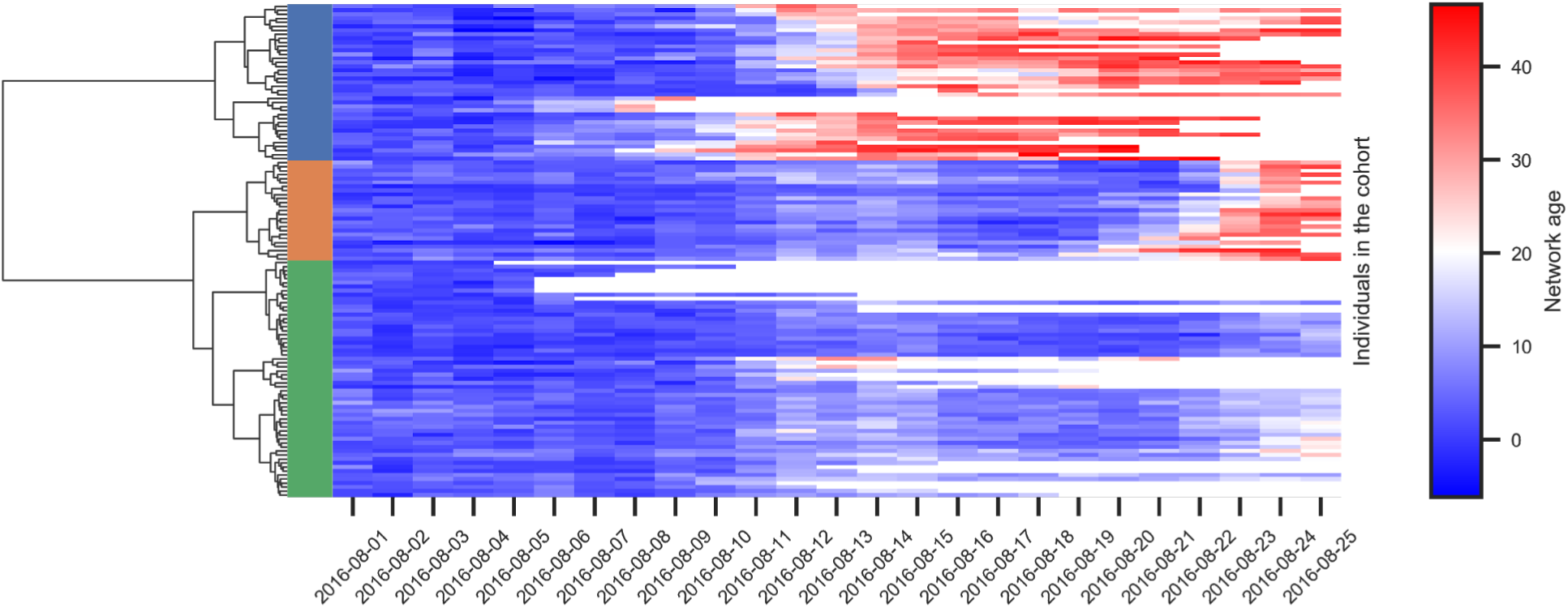
Agglomerative clustering of the network age over time of one cohort of bees (emerged on 2016-08-01) Missing values (e.g. due to the death of the bee) are white.

### SI 3.3 Threshold for bimodal network age distribution

For the bees depicted in fig. 3A, we calculate a threshold T=23.07 on the network age using Otsu’s method (Otsu 1979). The upper line contains all bees that fall above this threshold (N=832), the lower contains all bees below that threshold (N=563). The shaded areas depict 20%, 40%, 60% data intervals.

### SI 3.4 Quantifying when bees first split into distinct network age modes

We used KMeans to cluster the network age distribution of every day into two distinct clusters corresponding to the two modes. Then we check for every bee that we observed at least once as a young bee below the age of 6 (N=1079) at which age she first gets assigned to the higher cluster (mean=12.33, 95% CI = [6, 25.7], median=11, N=572). We ignore bees that are never assigned to the upper cluster (e.g. because we do not observe them for a long enough timespan). See fig. SI 15 for the distribution of biological ages.

### SI 3.5 Change of network age mode over a week

We analyze how many bees that are assigned to either of the network age modes on day X are assigned to the same cluster on day X+7 (clusters calculated as in section SI 3.4). Most of the bees in the upper cluster are also in the upper cluster one week later (mean=93% 95% CI [77%, 100%], N=3698 bee days).

Bees in the lower cluster tend to stay there less (mean=56% 95% CI [0%, 96%], N=10 927 bee days) depending on their age (see fig. SI 16).

**Figure SI 13:**
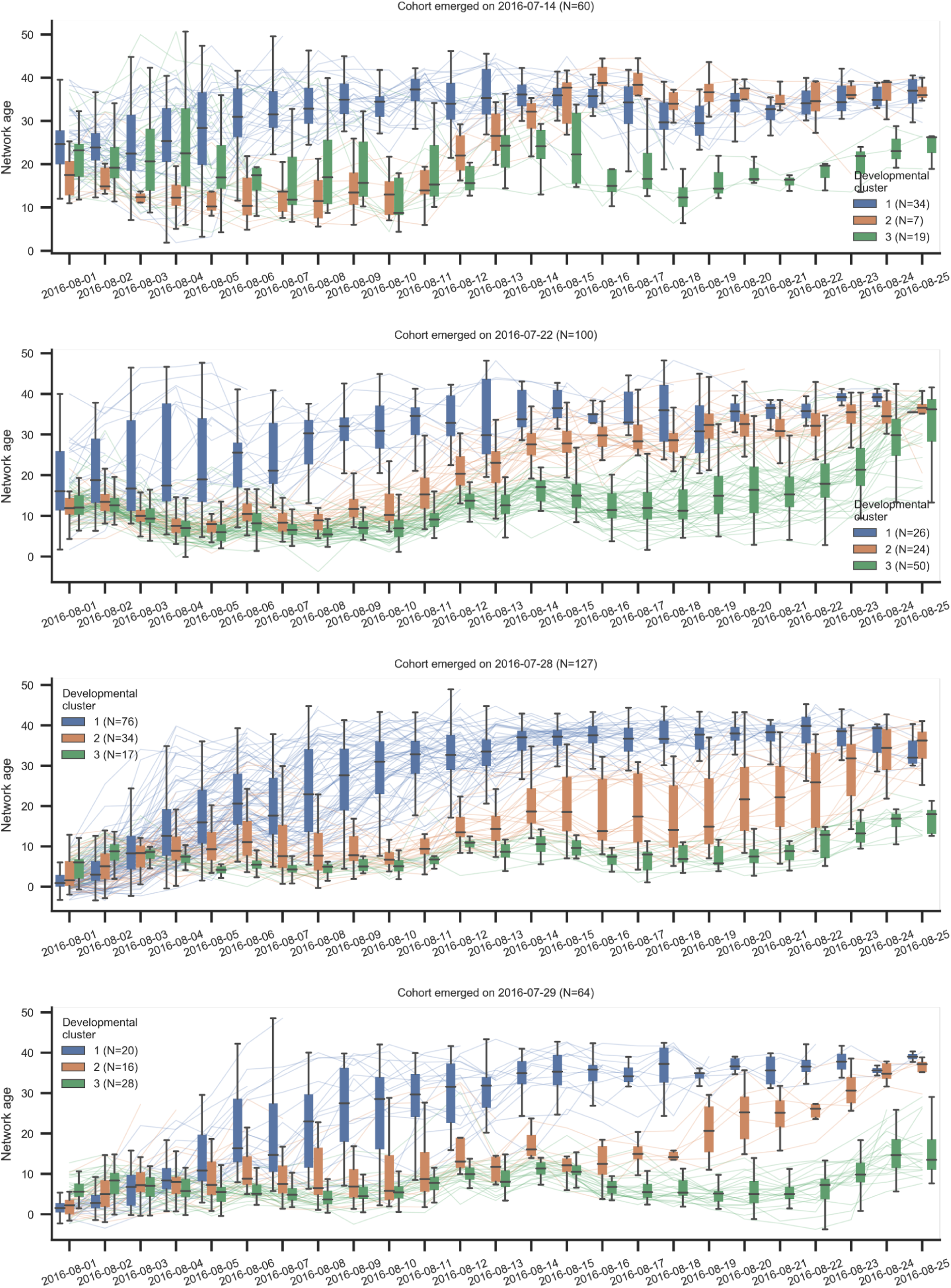
The development of the network age of different cohorts over the focal period (see individual titles). The individuals are grouped according to the agglomerative clustering. The lines depict the network age of individual bees. The boxplots summarize bees belonging to each cluster for a given day (center line, median; box limits, upper and lower quartiles; whiskers, 1.5x interquartile range).

**Figure SI 14:**
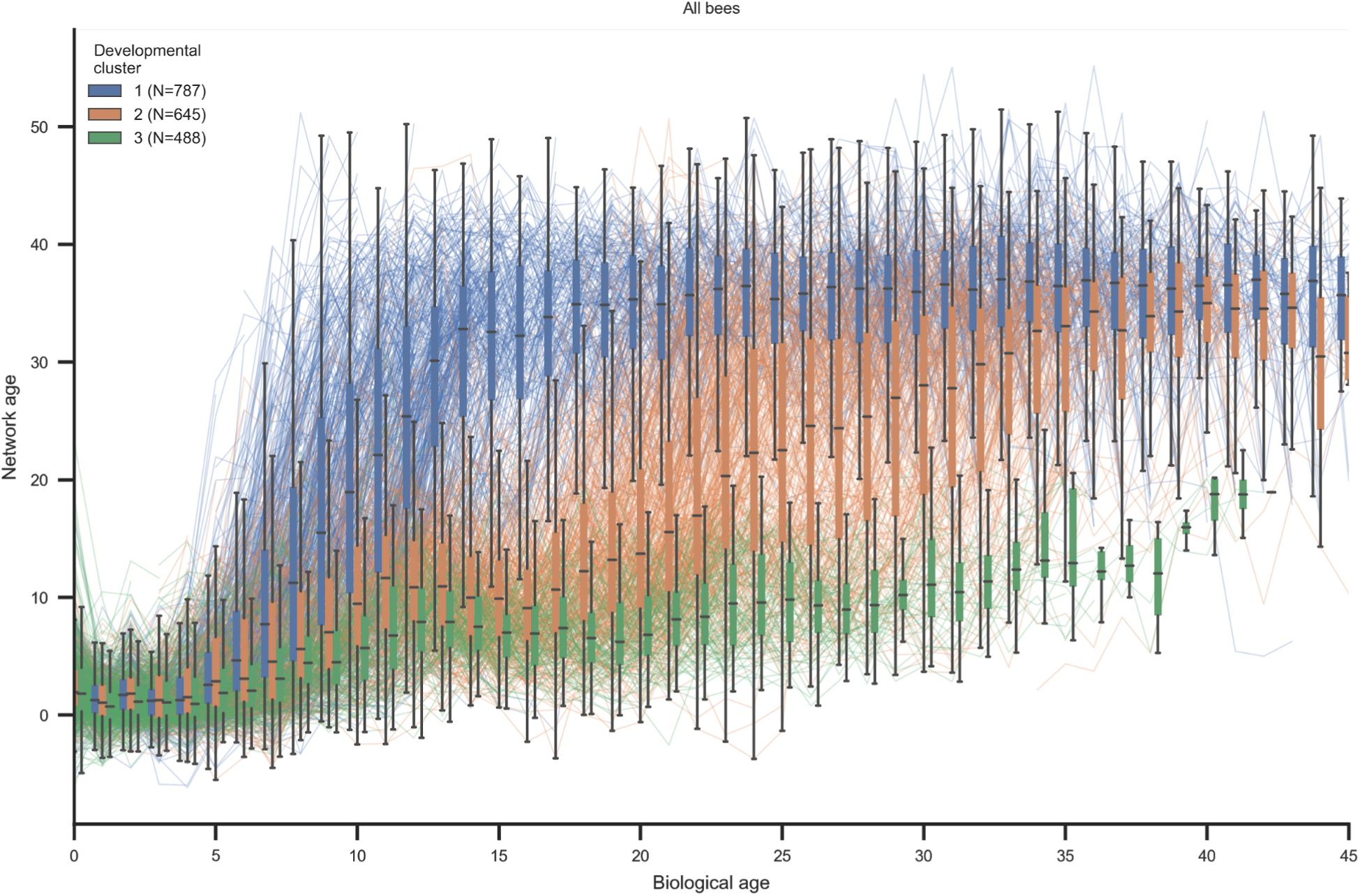
The network age development of all individuals over their age. To calculate this, we followed a procedure analogous to clustering the cohorts but used network age per age (instead of per date) as the bees’ feature vectors. Note that there was only a subset of 25 days of data available for each individual as some individuals were already in the hive as the focal period began and some were introduced later. The lines depict the network age of individual bees. The boxplots summarize bees belonging to each cluster for a given day (center line, median; box limits, upper and lower quartiles; whiskers, 1.5x interquartile range).

### SI 4 Network age predicts an individual’s behavior and future role in the colony

#### SI 4.1 Definition of circadian rhythmicity

The motion velocity of a bee was determined by dividing the euclidean distance between two consecutive detections by the time passed (a multiple of 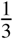 seconds). Any duplicate IDs were discarded. The velocity was median filtered with a kernel size of 3 to remove outliers. See fig. SI 17 for an example of the velocity of a specific bee over multiple days.

Lomb-Scargle periodograms were computed for all individuals remaining in the dataset at any time in the period from 2016-07-20 to 2016-09-18. For each day and individual, the Lomb-Scargle periodogram was calculated on the motion velocities over an interval of three days, i.e. including the preceding and following day. The circadian activity was confirmed as strong peaks at a period of 1 day. Lomb-Scargle periodograms were computed using the Astropy package (Collaboration et al. 2018). See fig. SI 17 for an example of a bee’s velocity and the resulting Lomb-Scargle periodogram.

In the following analyses, we reduced computational load by fitting a single sine wave of fixed frequency= 1/d (least-squares fit). For each fit, we extract the power as *P*(*f*) = 1 − (*S S E*_*sine*_*/S S E*_*constant*_) with *S S E*_*sine*_ being the residuals (sum of squared errors) of the sine fit and *S S E*_*constant*_ being the residuals (sum of squared errors) of a constant model assuming the mean of the data.

The power, hence, reflects how much of the velocity variation can be explained by the sinusoidal oscillation, or circadian rhythm.

**Figure SI 15:**
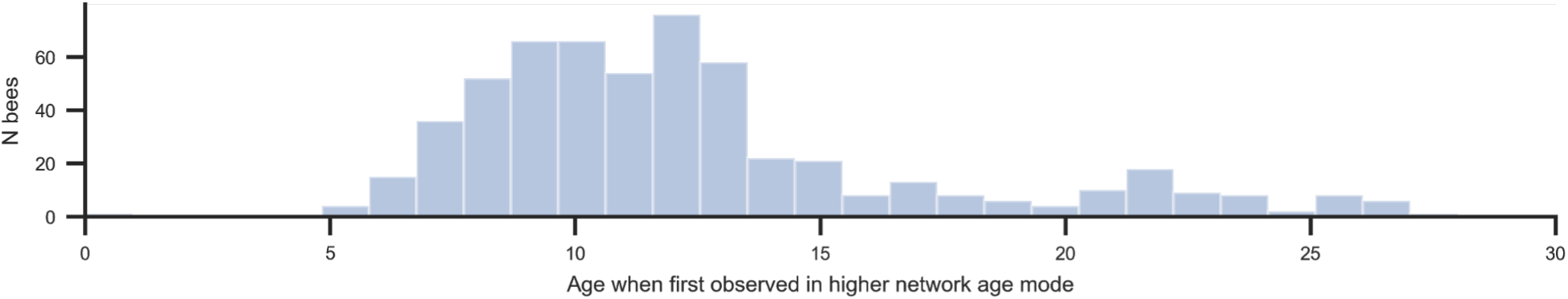
The distribution of the biological age at which a bee was first assigned to the higher network age cluster. Starting at around age 5-7, the network age distribution becomes bimodal.

**Figure SI 16:**
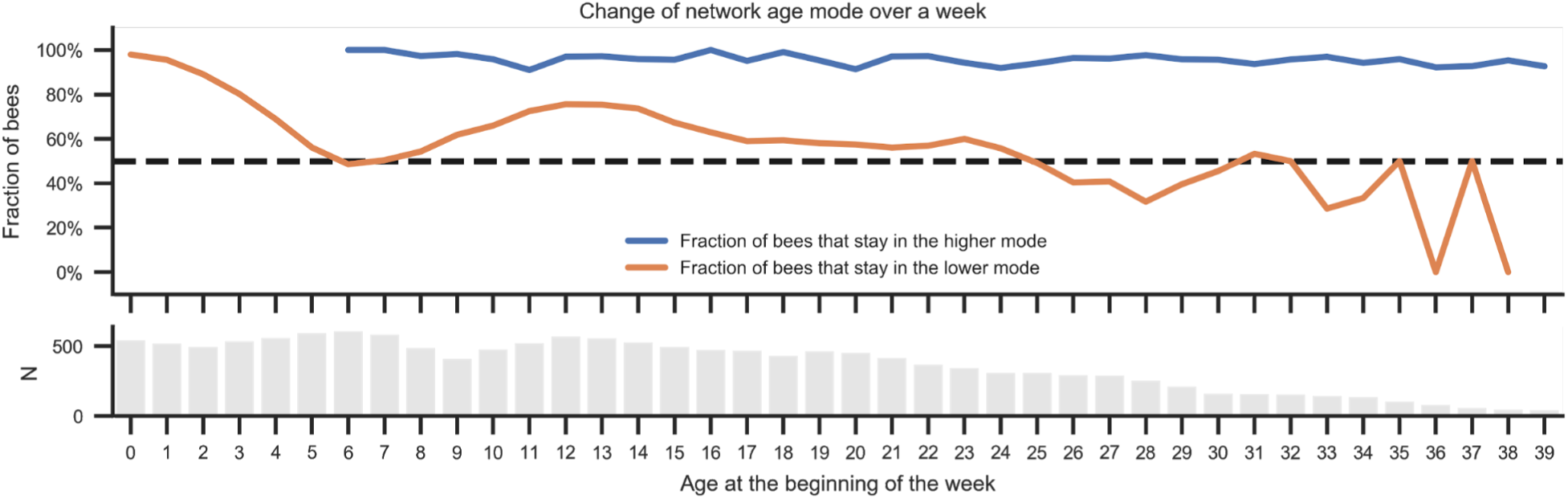
The consistency of network age cluster over the course of a week. (Top) We considered every individual and day (first day) for which we had data one week later (seventh day). We sorted them by their biological age on the first day (x-axis). Bees that were assigned to the higher network age cluster on the first day make up the blue line, bees from the lower cluster are depicted by the orange line. The y-axis value of each line is the fraction of bees that are still assigned to the same cluster seven days later. A high proportion of bees that are assigned to the higher cluster stay there over the course of one week (blue line). There are more bees that transition from the lower cluster to the higher one, regardless of age (orange line). (Bottom) The number of data points (orange and blue combined) for each age.

### SI 4.2 Targeted embedding using CCA

We show that we can extract targeted embeddings from the spectral factors of the interaction networks that are better in predicting other properties of the individuals. To compute those targeted embeddings, we exactly follow the methodology outlined in section SI 1.5, but for each property (days until death, time of peak activity, circadian rhythm, day- and nighttime velocities) we extract a one-dimensional embedding from the spectral factors that maximizes the correlation with this property using canonical-correlation analysis.

#### SI 4.3 Prediction of other behavior-related measures

We follow the same methodology as described in section SI 2.4 to evaluate how well the various age measures explain various properties of the individuals with a slight adjustment: We choose the identity function as the link function of the generalized linear model and as the activation function of the neural network’s final layer. We model the residual distribution as a normal distribution with constant variance.

**Figure SI 17:**
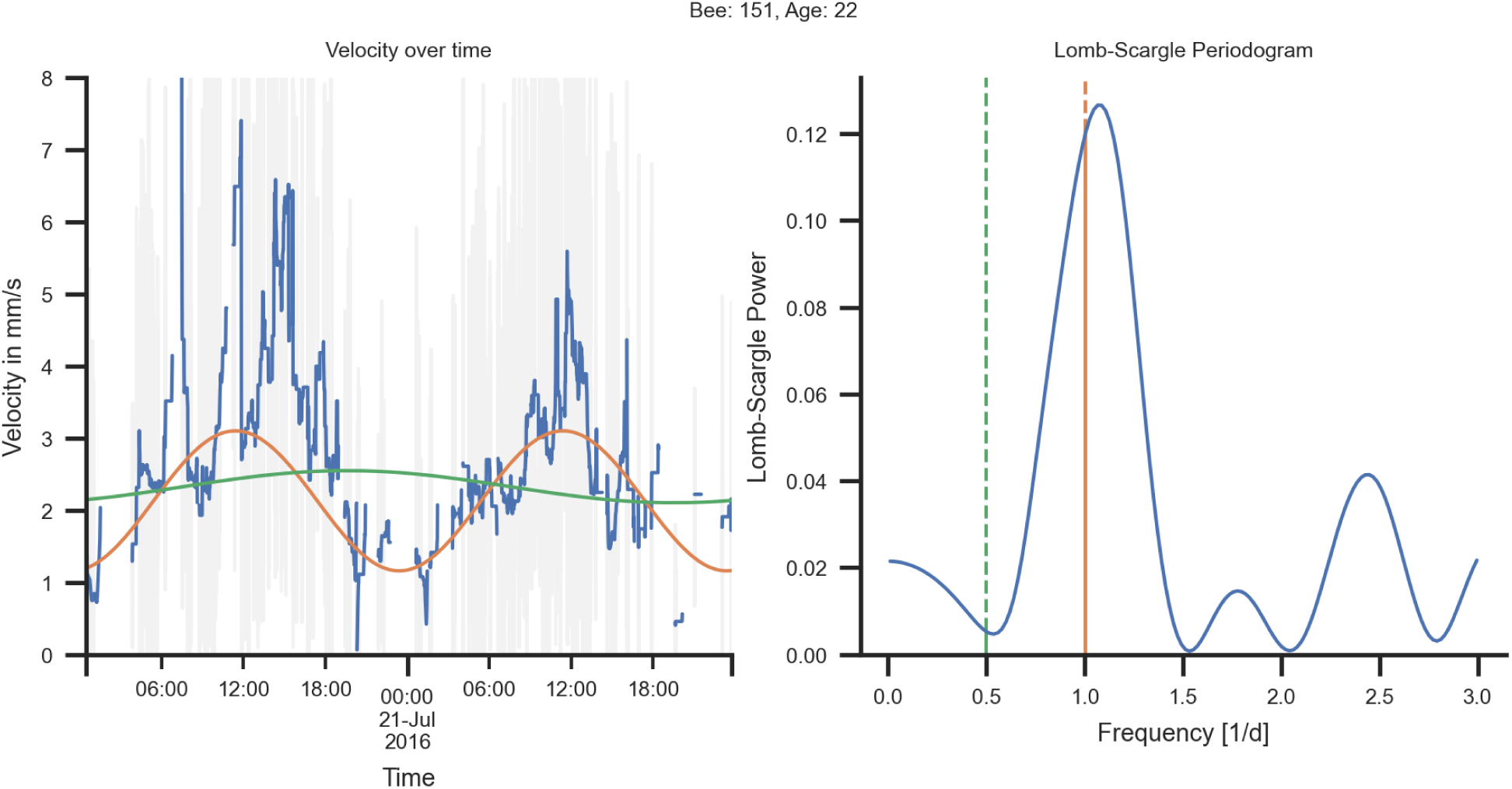
Calculation of circadian rhythmicity from movement data. (Left) The velocity as raw data (grey line) and smoothed with a median filter (blue line) for one bee over two days. The orange sine wave has a period of one day and was fitted to the velocity data via least-squares fitting. The green line is a similar fit with a period of two days. (Right) The corresponding Lomb-Scargle periodogram with the powers at the one-day frequency (orange) and two-day frequencies (green) highlighted.

We define the ‘days until death’ as the number of days left in an individual’s life on a given day (for a description of the automatically determined death dates see section SI 1.2). The time of peak activity and the rhythmicity of daily movement were calculated as described in section SI 4.1.

For the daytime and nighttime velocities we use the same data as for the circadian rhythmicity (see section SI 4.1). For the daytime velocities we use the mean of all collected velocities between 09:00-18:00 UTC of the three-day rolling window; for nighttime velocities, 21:00-06:00 UTC.

See table 4a for an overview of the scores of different models and targets.

### SI 4.4 Future predictability

We evaluate how well we can predict future task allocation using network age and biological age. To ensure that no information leak can occur, we only used supervised information from the past and test it on future data. To do this, we first calculate the spectral factors for the entire dataset as described in section SI 1.5. The factors are computed for each day separately, hence no information can leak from the past to the future. We then determine the mapping from spectral factors to network age using CCA, but only on a fixed range of days prior to the validation window. We fix the number of days in the train set to 12 days so that we always have approximately the same amount of training data independent of the number of days we predict into the future. Similar to the linear mapping given by CCA, we also determine the parameters of the regression model only on the train dataset from the same fixed time window. We train separate models for all viable ranges of dates and for prediction from one to 11 days into the future. The linear mapping given by the CCA and the predictive models are then applied to the spectral factors on the held-out validation set from a time after the training data set (see fig. SI 20 for an overview about the data handling). For this analysis, we want to evaluate how well we can predict task allocation into the future. We estimate the effect size in 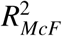 by calculating the 95% CI for the different time windows (see fig. SI 20, median improvement in 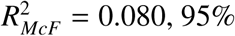 CI [0.055, 0.090], N=12).

**Table 4:**
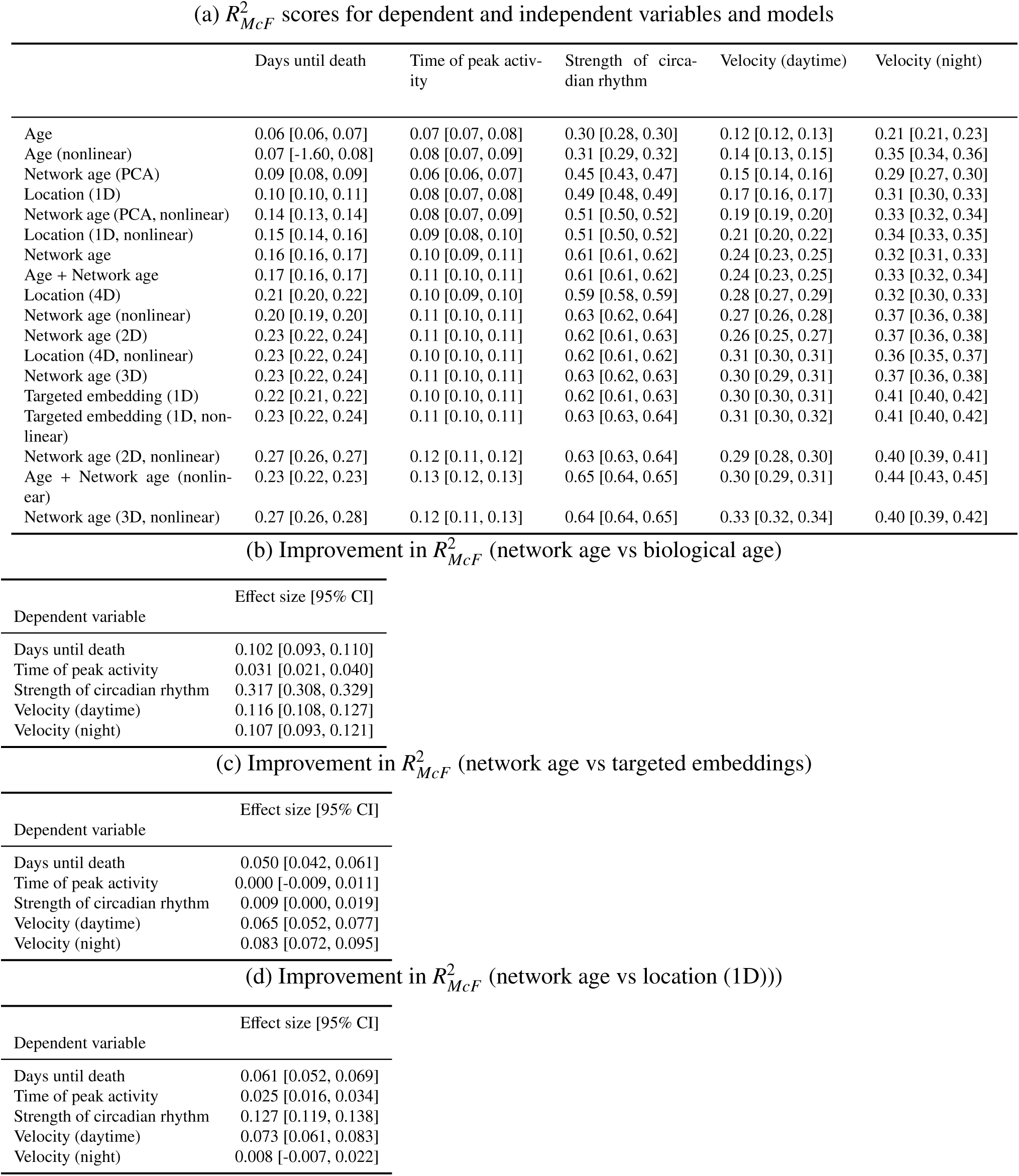
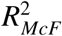 values for different models when predicting different behavior-related metrics. Biological age consistently performs worse than predictions based on network age. Our method can also be applied without spatial information by optimizing the CCA for these metrics, yielding embeddings that are strongly predictive of them while still only using information available from the social networks (see *Targeted embedding (1D)*). (Bottom) 95% bootstrap confidence intervals of the effect sizes (b) *R*^2^ values for network age and biological age, c) 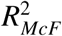 values for targeted embeddings and network age, N=128).

|  | Days until death | Time of peak activity | Strength of circadian rhythm | Velocity (daytime) | Velocity (night) |
| --- | --- | --- | --- | --- | --- |
| Age | 0.06 [0.06, 0.07] | 0.07 [0.07, 0.08] | 0.30 [0.28, 0.30] | 0.12 [0.12, 0.13] | 0.21 [0.21, 0.23] |
| Age (nonlinear) | 0.07 [-1.60, 0.08] | 0.08 [0.07, 0.09] | 0.31 [0.29, 0.32] | 0.14 [0.13, 0.15] | 0.35 [0.34, 0.36] |
| Network age (PCA) | 0.09 [0.08, 0.09] | 0.06 [0.06, 0.07] | 0.45 [0.43, 0.47] | 0.15 [0.14, 0.16] | 0.29 [0.27, 0.30] |
| Location (1D) | 0.10 [0.10, 0.11] | 0.08 [0.07, 0.08] | 0.49 [0.48, 0.49] | 0.17 [0.16, 0.17] | 0.31 [0.30, 0.33] |
| Network age (PCA, nonlinear) | 0.14 [0.13, 0.14] | 0.08 [0.07, 0.09] | 0.51 [0.50, 0.52] | 0.19 [0.19, 0.20] | 0.33 [0.32, 0.34] |
| Location (1D, nonlinear) | 0.15 [0.14, 0.16] | 0.09 [0.08, 0.10] | 0.51 [0.50, 0.52] | 0.21 [0.20, 0.22] | 0.34 [0.33, 0.35] |
| Network age | 0.16 [0.16, 0.17] | 0.10 [0.09, 0.11] | 0.61 [0.61, 0.62] | 0.24 [0.23, 0.25] | 0.32 [0.31, 0.33] |
| Age + Network age | 0.17 [0.16, 0.17] | 0.11 [0.10, 0.11] | 0.61 [0.61, 0.62] | 0.24 [0.23, 0.25] | 0.33 [0.32, 0.34] |
| Location (4D) | 0.21 [0.20, 0.22] | 0.10 [0.09, 0.10] | 0.59 [0.58, 0.59] | 0.28 [0.27, 0.29] | 0.32 [0.30, 0.33] |
| Network age (nonlinear) | 0.20 [0.19, 0.20] | 0.11 [0.10, 0.11] | 0.63 [0.62, 0.64] | 0.27 [0.26, 0.28] | 0.37 [0.36, 0.38] |
| Network age (2D) | 0.23 [0.22, 0.24] | 0.11 [0.10, 0.11] | 0.62 [0.61, 0.63] | 0.26 [0.25, 0.27] | 0.37 [0.36, 0.38] |
| Location (4D, nonlinear) | 0.23 [0.22, 0.24] | 0.10 [0.10, 0.11] | 0.62 [0.61, 0.62] | 0.31 [0.30, 0.31] | 0.36 [0.35, 0.37] |
| Network age (3D) | 0.23 [0.22, 0.24] | 0.11 [0.10, 0.11] | 0.63 [0.62, 0.63] | 0.30 [0.29, 0.31] | 0.37 [0.36, 0.38] |
| Targeted embedding (1D) | 0.22 [0.21, 0.22] | 0.10 [0.10, 0.11] | 0.62 [0.61, 0.63] | 0.30 [0.30, 0.31] | 0.41 [0.40, 0.42] |
| Targeted embedding (1D, nonlinear) | 0.23 [0.22, 0.24] | 0.11 [0.10, 0.11] | 0.63 [0.63, 0.64] | 0.31 [0.30, 0.32] | 0.41 [0.40, 0.42] |
| Network age (2D, nonlinear) | 0.27 [0.26, 0.27] | 0.12 [0.11, 0.12] | 0.63 [0.63, 0.64] | 0.29 [0.28, 0.30] | 0.40 [0.39, 0.41] |
| Age + Network age (nonlinear) | 0.23 [0.22, 0.23] | 0.13 [0.12, 0.13] | 0.65 [0.64, 0.65] | 0.30 [0.29, 0.31] | 0.44 [0.43, 0.45] |
| Network age (3D, nonlinear) | 0.27 [0.26, 0.28] | 0.12 [0.11, 0.13] | 0.64 [0.64, 0.65] | 0.33 [0.32, 0.34] | 0.40 [0.39, 0.42] |

(b) Improvement in $R^2_{McF}$ (network age vs biological age)
| Dependent variable | Effect size [95% CI] |
| --- | --- |
| Days until death | 0.102 [0.093, 0.110] |
| Time of peak activity | 0.031 [0.021, 0.040] |
| Strength of circadian rhythm | 0.317 [0.308, 0.329] |
| Velocity (daytime) | 0.116 [0.108, 0.127] |
| Velocity (night) | 0.107 [0.093, 0.121] |

(c) Improvement in $R^2_{McF}$ (network age vs targeted embeddings)
| Dependent variable | Effect size [95% CI] |
| --- | --- |
| Days until death | 0.050 [0.042, 0.061] |
| Time of peak activity | 0.000 [-0.009, 0.011] |
| Strength of circadian rhythm | 0.009 [0.000, 0.019] |
| Velocity (daytime) | 0.065 [0.052, 0.077] |
| Velocity (night) | 0.083 [0.072, 0.095] |

Table 4: $R^2_{McF}$ values for different models when predicting different behavior-related metrics.
| Dependent variable | Effect size [95% CI] |
| --- | --- |
| Days until death | 0.061 [0.052, 0.069] |
| Time of peak activity | 0.025 [0.016, 0.034] |
| Strength of circadian rhythm | 0.127 [0.119, 0.138] |
| Velocity (daytime) | 0.073 [0.061, 0.083] |
| Velocity (night) | 0.008 [-0.007, 0.022] |

**Figure SI 18:**
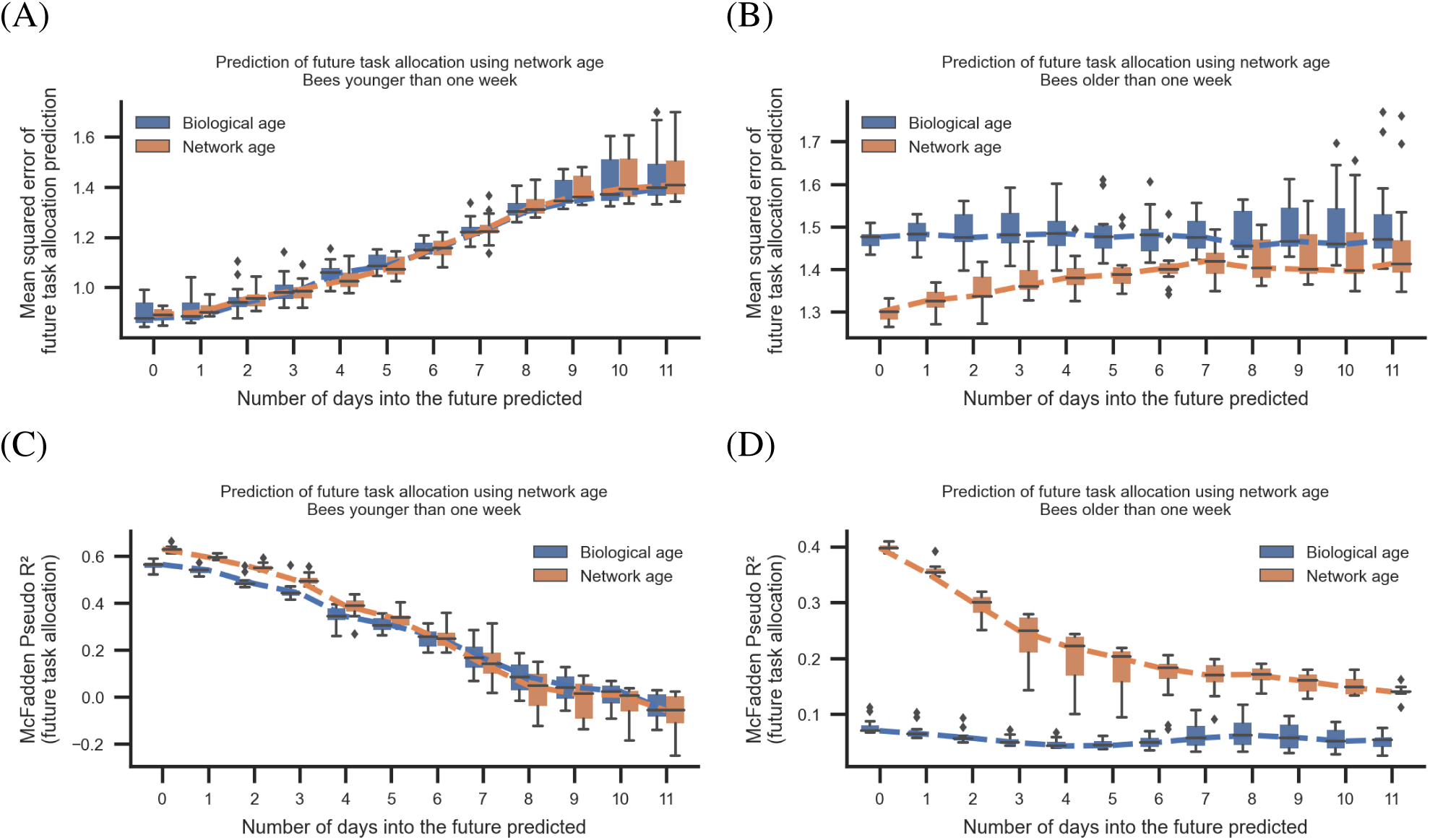
Mean squared error (top row) and McFadden’s *R*^2^ (bottom row) for a model that predicts task allocation up to 11 days into the future. We distinguish between young bees (left column) and old bees (right column). For the old bees (> 7 days of age), network age consistently performs better than age. When predicting task allocation 11 days into the future for old bees, network age performs better than age performs for the current date. For young bees, we find neither biological age nor network age to be predictive. Each box comprises 128 bootstrap runs (center line, median; box limits, upper and lower quartiles; whiskers, 1.5x interquartile range; points, outliers).

Because the two compared models are not nested, the likelihood ratio test does not apply here. We perform a paired binomial test using the null hypothesis that the improvement in mean squared error in task allocation prediction is zero or less. We find that we can predict the task allocation of an individual seven days into the future with a lower mean squared error (paired binomial test, *p* 0.001, N=55 390) using network age.

This is mostly caused by older bees. For young bees, there is a low amount of variance in task allocation and network age is about as good in describing task allocation for young bees as biological age. Furthermore, we find that we can not reliably predict future task allocation for young bees, suggesting that either the social networks in this study are not predictive for this task or that the future task allocation of young bees is driven by other factors. See fig. SI 18 for an overview of the results.

The predictive power of network age cannot be fully explained by the bimodal distribution of network age and organization of stable *work groups* within the colony. We perform an additional analysis to reject this repeatability null hypothesis by comparing how well the task allocation prediction using networks works when simply shifting the prediction for the current day into the future. We find that a model fitted to predict future task allocation outperforms this null model considerably (See fig. SI 19).

**Figure SI 19:**
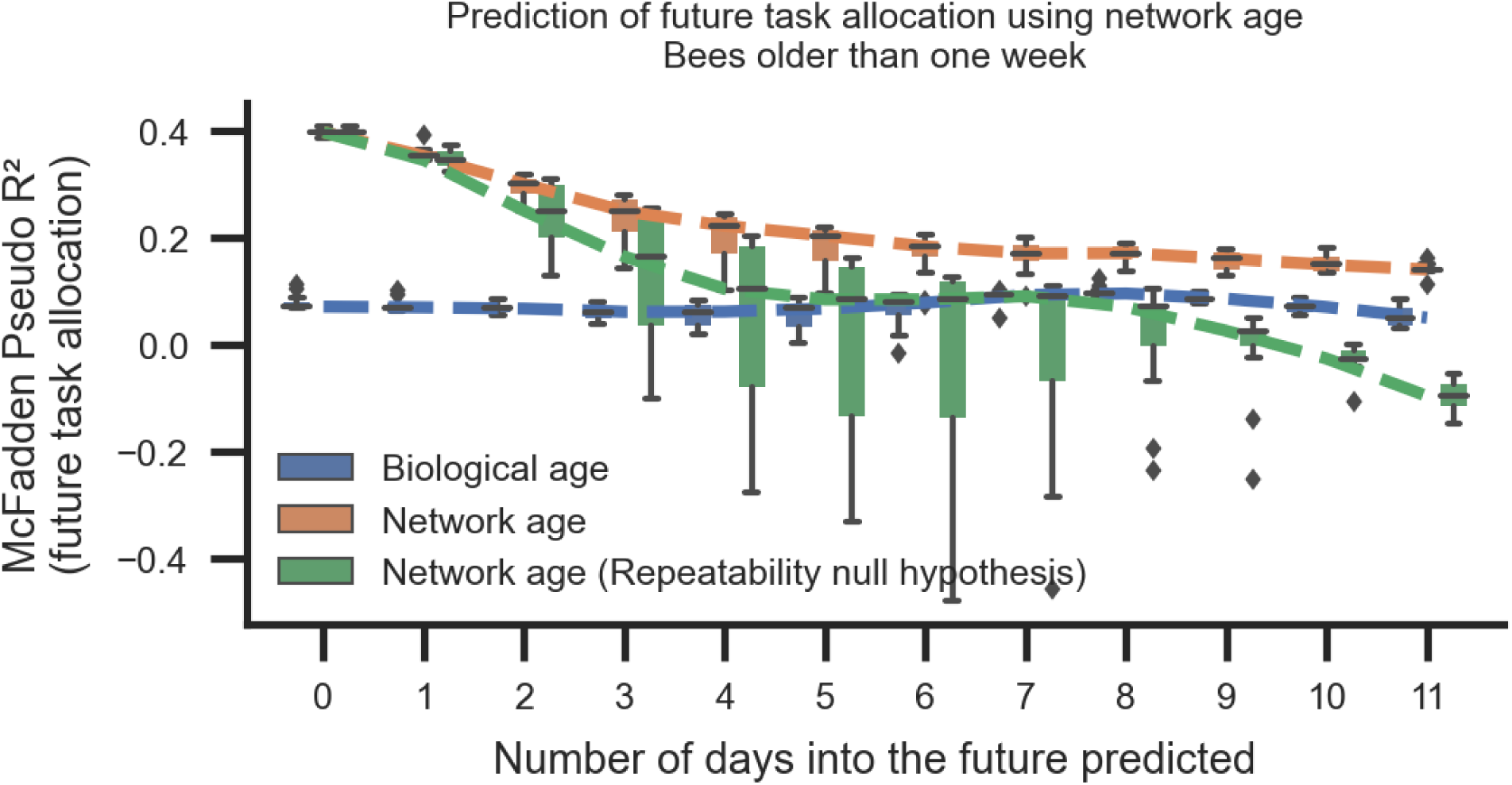
Ruling out repeatability as a cause for predictiveness of network age. To check that the predictiveness of network age for old bees is not simply due to the bimodal distribution of the network age and its high repeatability, we compare a predictive model based on network age (orange) with the null-hypothesis that the network age of day X is sufficient to describe the coming days without explicitly modeling the changes. Each box comprises 128 bootstrap runs (center line, median; box limits, upper and lower quartiles; whiskers, 1.5x interquartile range; points, outliers).

### SI 4.5 Probability of dying over the course of a week

We have shown that network age is more predictive of the remaining days to live than biological age (see section SI 4.3). Here we directly compare biologically young but functionally old bees with biologically old but functionally young bees. To define biologically young and old, we orientate ourselves at the caste thresholds proposed by Seeley (Seeley 1982). To define functionally young and old we use the same thresholds for network age as shown in the heatmaps of fig. 3A, because they occupy distinct locations on the comb and therefore hold different functional roles within the colony. However, these exact thresholds are not important as both the effect strength and the significance are high.

We sort the individuals into the two groups based on their network age and biological age. To not count an individual twice, we only consider the last date that a bee was seen in each group. We calculate the probability of dying P as the fraction of bees of each group we do not consider alive after one week (see section SI 1.2 for more information about the death date of a bee). We calculate a confidence interval by drawing 1000 bootstrap samples for each group.

Biologically young but functionally old bees (biological age < 11 days, network age > 30) have a probability of dying over the course of a week of P=80.6% (bootstrap 95% CI=[74.1%, 87.1%], N=139). Biologically old but functionally young bees (biological age > 20, network age < 15) have a significantly lower probability of dying of P=42.1% (bootstrap 95% CI=[37.4%, 47.2%], N=390; *χ*^2^ test of independence *p* ≪ 0.001 N=529).

**Figure SI 20:**
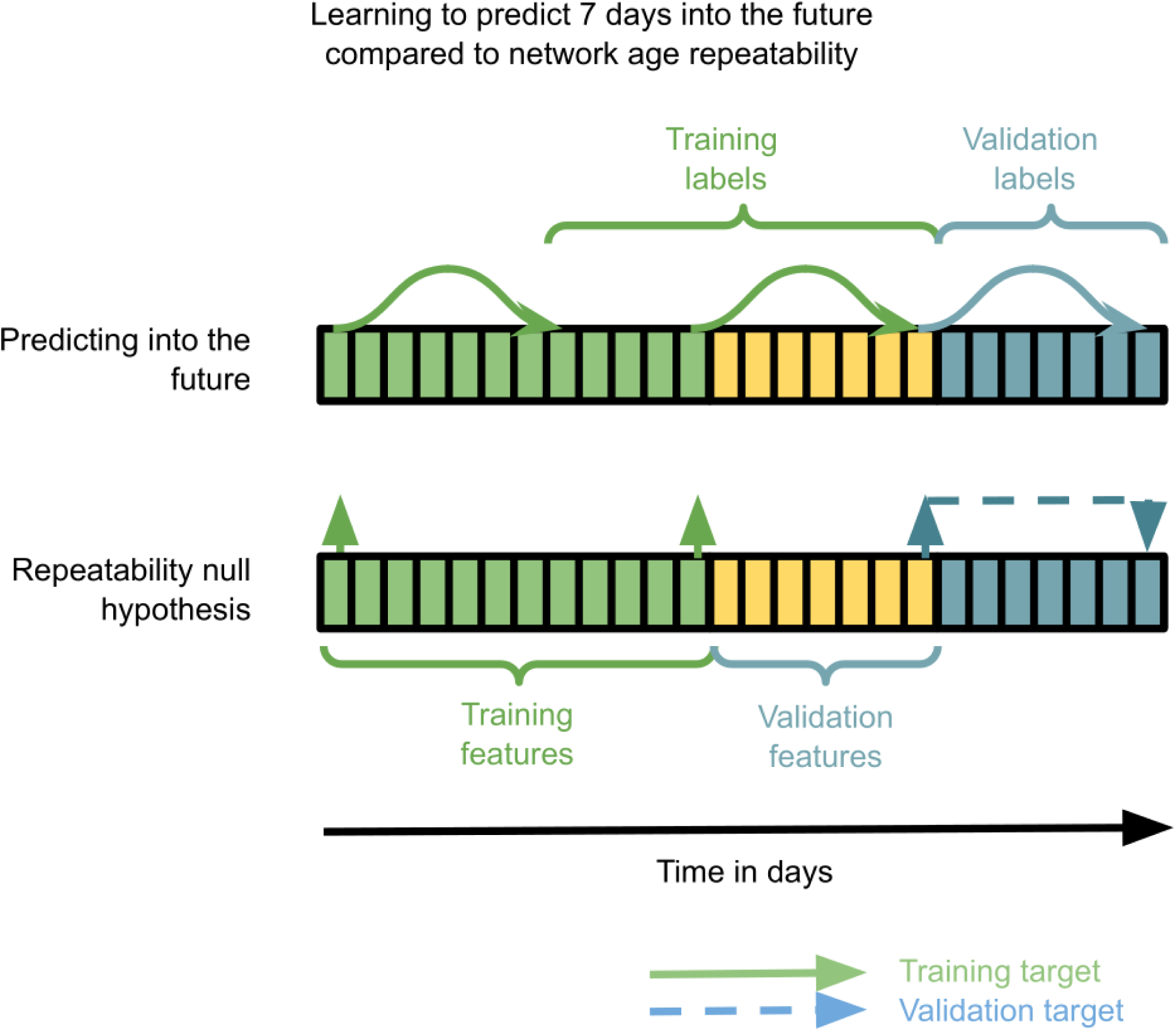
Data handling for cross-validating future predictability. (Top) Depiction of the process of training a model based on either biological age or network age to predict the future task allocation. We make sure that we do not leak information when evaluating future predictiveness. (Bottom) Data handling for the null hypothesis to check that the predictiveness is not completely explained by the repeatability of network age and task allocation for individual bees.

### SI 4.6 Time spent in task-associated locations as a predictor

To evaluate how well the time spent in the measured task-associated locations itself is able to predict an individual’s behavior and future role in the colony, we derive two representations of this spatial information and use them as predictors in the same way as network age (as described in sections SI 4.3 and SI 4.4) for predicting mortality and movement patterns.

We use the task descriptor described in section SI 1.4 either directly as the independent variables in the regression models (*Location (4D)* in table 4a) or a 1D representation of this descriptor derived using PCA (*Location (1D)* in table 4a). The 4D descriptor is not directly comparable to network age because of the higher dimensionality, but serves as an upper bound on the information contained in the task descriptor. In terms of dimensionality, the 1D representation can directly be compared to network age.

We find that this spatial information is a better descriptor of an individual’s behavior than age, but significantly less informative when compared to network age (table 4d). See table 4 for an overview of the scores and effect sizes.

## References

Abdi, Hervé (2007). “Singular and Generalized Singular Value Decomposition”. In: Encyclopedia of Measurement and Statistics. 3 vols. Thousand Oaks: SAGE Publications, Inc., pp. 908–912. doi: 10.4135/9781412952644.

Amdam, Gro V., Kari Norberg, et al. (Aug. 3, 2004). “Reproductive Ground Plan May Mediate Colony-Level Selection Effects on Individual Foraging Behavior in Honey Bees”. In: Proceedings of the National Academy of Sciences 101.31, pp. 11350–11355. doi: 10.1073/pnas.0403073101.

Amdam, Gro V. and Robert E. Page Jr (2010). “The Developmental Genetics and Physiology of Honeybee Societies”. In: Animal behaviour 79.5, pp. 973–980.

Amdam, Gro Vang and Stig W. Omholt (Aug. 2003). “The Hive Bee to Forager Transition in Honeybee Colonies: The Double Repressor Hypothesis”. In: Journal of Theoretical Biology 223.4, pp. 451–464. doi: 10.1016/S0022-5193(03)00121-8.

Ament, Seth A., Ying Wang, and Gene E. Robinson (2010). “Nutritional Regulation of Division of Labor in Honey Bees: Toward a Systems Biology Perspective”. In: WIREs Systems Biology and Medicine 2.5, pp. 566–576. doi: 10.1002/wsbm.73.

Aplin, Lucy M. et al. (Feb. 26, 2015). “Experimentally Induced Innovations Lead to Persistent Culture via Conformity in Wild Birds”. In: Nature 518.7540, pp. 538–541. doi: 10.1038/nature13998.

Balbuena, M. S., J. Molinas, and W. M. Farina (Mar. 1, 2012). “Honeybee Recruitment to Scented Food Sources: Correlations between in-Hive Social Interactions and Foraging Decisions”. In: Behavioral Ecology and Sociobiology 66.3, pp. 445–452. doi: 10.1007/s00265-011-1290-3.

Baracchi, David and Alessandro Cini (2014). “A Socio-Spatial Combined Approach Confirms a Highly Compartmentalised Structure in Honeybees”. In: Ethology 120.12, pp. 1167–1176. doi: 10.1111/eth.12290.

Belkin, Mikhail and Partha Niyogi (June 2003). “Laplacian Eigenmaps for Dimensionality Reduction and Data Representation”. In: Neural Computation 15.6, pp. 1373–1396. doi: 10.1162/089976603321780317.

Blut, Christina et al. (Dec. 15, 2017). “Automated Computer-Based Detection of Encounter Behaviours in Groups of Honeybees”. In: Scientific Reports 7.1, p. 17663. doi: 10.1038/s41598-017-17863-4.

Boenisch, Franziska et al. (2018). “Tracking All Members of a Honey Bee Colony Over Their Lifetime Using Learned Models of Correspondence”. In: Frontiers in Robotics and AI 5, p. 35.

Bozek, Katarzyna et al. (Mar. 27, 2020). “Markerless Tracking of an Entire Insect Colony”. In: bioRxiv, p. 2020.03.26.007302. doi: 10.1101/2020.03.26.007302.

Cheney, Dorothy L., Joan B. Silk, and Robert M. Seyfarth (2016). “Network Connections, Dyadic Bonds and Fitness in Wild Female Baboons”. In: Royal Society Open Science 3.7, p. 160255. doi: 10.1098/rsos.160255.

Cholé, Hanna et al. (Apr. 22, 2019). “Social Contact Acts as Appetitive Reinforcement and Supports Associative Learning in Honeybees”. In: Current Biology 29.8, 1407–1413.e3. doi: 10.1016/j.cub.2019.03.025.

Collaboration, The Astropy et al. (Aug. 23, 2018). “The Astropy Project: Building an Inclusive, Open-Science Project and Status of the v2.0 Core Package”. In: The Astronomical Journal 156.3, p. 123. doi: 10.3847/1538-3881/aabc4f.

Crall, James D. et al. (Apr. 3, 2018). “Spatial Fidelity of Workers Predicts Collective Response to Disturbance in a Social Insect”. In: Nature Communications 9.1, p. 1201. doi: 10.1038/s41467-018-03561-w.

Davidson, Jacob D. and Deborah M. Gordon (Oct. 31, 2017). “Spatial Organization and Interactions of Harvester Ants during Foraging Activity”. In: Journal of The Royal Society Interface 14.135, p. 20170413. doi: 10.1098/rsif.2017.0413.

Dreller, Claudia, Robert E. Page Jr, and M. Kim Fondrk (Mar. 1, 1999). “Regulation of Pollen Foraging in Honeybee Colonies: Effects of Young Brood, Stored Pollen, and Empty Space”. In: Behavioral Ecology and Sociobiology 45.3-4, pp. 227–233. doi: 10.1007/s002650050557.

Farina, Walter Marcelo (2000). “The Interplay between Dancing and Trophallactic Behavior in the Honey Bee Apis Mellifera”. In: Journal of Comparative Physiology A. doi: 10.1007/s003590050424.

Farine, Damien R. and Hal Whitehead (2015). “Constructing, Conducting and Interpreting Animal Social Network Analysis”. In: Journal of Animal Ecology 84.5, pp. 1144–1163.

Flack, Jessica C. et al. (Jan. 2006). “Policing Stabilizes Construction of Social Niches in Primates”. In: Nature 439.7075, pp. 426–429. doi: 10.1038/nature04326.

Frisch, Karl von (1967). “The Dance Language and Orientation of Bees.” In:

Geffre, Amy C. et al. (Apr. 22, 2020). “Honey Bee Virus Causes Context-Dependent Changes in Host Social Behavior”. In: Proceedings of the National Academy of Sciences. doi: 10.1073/pnas.2002268117.

Gernat, Tim et al. (Feb. 13, 2018). “Automated Monitoring of Behavior Reveals Bursty Interaction Patterns and Rapid Spreading Dynamics in Honeybee Social Networks”. In: Proceedings of the National Academy of Sciences 115.7, pp. 1433–1438. doi: 10.1073/pnas.1713568115.

Girard, M. B., H. R. Mattila, and T. D. Seeley (Feb. 1, 2011). “Recruitment-Dance Signals Draw Larger Audiences When Honey Bee Colonies Have Multiple Patrilines”. In: Insectes Sociaux 58.1, pp. 77–86. doi: 10.1007/s00040-010-0118-x.

Gomez-Marin, Alex et al. (Nov. 1, 2014). “Big Behavioral Data: Psychology, Ethology and the Foundations of Neuroscience”. In: Nature Neuroscience 17.11, pp. 1455–1463.

Gordon, Deborah M. (Mar. 1996). “The Organization of Work in Social Insect Colonies”. In: Nature 380.6570 (6570), pp. 121–124. doi: 10.1038/380121a0.

Gordon, Deborah M. (2010). Ant Encounters: Interaction Networks and Colony Behavior. Princeton University Press.

Goyret, J. and W. M. Farina (Aug. 1, 2003). “Descriptive Study of Antennation during Trophallactic Unloading Contacts in Honeybees Apis Mellifera Carnica”. In: Insectes Sociaux 50.3, pp. 274–276. doi: 10.1007/s00040-003-0678-0.

Hasenjager, Matthew J., William Hoppitt, and Ellouise Leadbeater (Jan. 31, 2020). “Network-Based Diffusion Analysis Reveals Context-Specific Dominance of Dance Communication in Foraging Honeybees”. In: Nature Communications 11.1 (1), pp. 1–9. doi: 10.1038/s41467-020-14410-0.

Hotelling, Harold (Dec. 1, 1936). “Relations between Two Sets of Variables”. In: Biometrika 28.3-4, pp. 321–377. doi: 10.1093/biomet/28.3-4.321.

Huang, Z. Y. and G. E. Robinson (Apr. 1, 1995). “Seasonal Changes in Juvenile Hormone Titers and Rates of Biosynthesis in Honey Bees”. In: Journal of Comparative Physiology B 165.1, pp. 18–28. doi: 10.1007/BF00264682.

Huang, Z. Y. (Sept. 1, 1996). “Regulation of Honey Bee Division of Labor by Colony Age Demography”. In: Behavioral Ecology and Sociobiology 39.3, pp. 147–158. doi: 10.1007/s002650050276.

Ihle, Kate E. et al. (May 1, 2010). “Genotype Effect on Regulation of Behaviour by Vitellogenin Supports Reproductive Origin of Honeybee Foraging Bias”. In: Animal Behaviour 79.5, pp. 1001–1006. doi: 10.1016/j.anbehav.2010.02.009.

Johnson, Brian R. (Jan. 2010). “Division of Labor in Honeybees: Form, Function, and Proximate Mechanisms”. In: Behavioral Ecology and Sociobiology 64.3, pp. 305–316. doi: 10.1007/s00265-009-0874-7.

Knapp, Thomas R. (1978). “Canonical Correlation Analysis: A General Parametric Significance-Testing System”. In: Psychological Bulletin 85.2, pp. 410–416. doi: 10.1037/0033-2909.85.2.410.

Krause, Jens et al. (2015). Animal Social Networks. Oxford University Press. 279 pp.

Levene, H. (1960). “Robust Tests for Equality of Variances, p 278–292”. In: Contributions to probability and statistics: essays in honor of Harold Hotelling. Stanford University Press, Palo Alto, CA.

Lusseau, David and M. E. J. Newman (2004). “Identifying the Role That Animals Play in Their Social Networks”. In: Proceedings: Biological Sciences 271, S477–S481.

McFadden, Daniel (1973). Conditional Logit Analysis of Qualitative Choice Behavior.

Mersch, Danielle P., Alessandro Crespi, and Laurent Keller (2013). “Tracking Individuals Shows Spatial Fidelity Is a Key Regulator of Ant Social Organization”. In: Science 340.6136, pp. 1090–1093.

Mönck, Hauke Jürgen (2016). “Large Scale Supercomputer Assisted and Live Video Encoding with Image Statistics”. Masters Thesis. Freie Universität Berlin.

Naug, Dhruba (Sept. 1, 2008). “Structure of the Social Network and Its Influence on Transmission Dynamics in a Honeybee Colony”. In: Behavioral Ecology and Sociobiology 62.11, pp. 1719–1725. doi: 10.1007/s00265-008-0600-x.

Nieh, James C. (Feb. 23, 2010). “A Negative Feedback Signal That Is Triggered by Peril Curbs Honey Bee Recruitment”. In: Current Biology 20.4, pp. 310–315. doi: 10.1016/j.cub.2009.12.060.

Otsu, Nobuyuki (Jan. 1979). “A Threshold Selection Method from Gray-Level Histograms”. In: IEEE Transactions on Systems, Man, and Cybernetics 9.1, pp. 62–66. doi: 10.1109/TSMC.1979.4310076.

Pankiw, T. and R. E. Page Jr. (Aug. 1, 1999). “The Effect of Genotype, Age, Sex, and Caste on Response Thresholds to Sucrose and Foraging Behavior of Honey Bees (Apis Mellifera L.)” In: Journal of Comparative Physiology A 185.2, pp. 207–213. doi: 10.1007/s003590050379.

Paszke, Adam et al. (2019). “PyTorch: An Imperative Style, High-Performance Deep Learning Library”. In: p. 12.

Pedregosa, Fabian et al. (2011). “Scikit-Learn: Machine Learning in Python”. In: Journal of Machine Learning Research 12 (Oct), pp. 2825–2830.

Perry, Clint J. et al. (Mar. 17, 2015). “Rapid Behavioral Maturation Accelerates Failure of Stressed Honey Bee Colonies”. In: Proceedings of the National Academy of Sciences 112.11, pp. 3427–3432. doi: 10.1073/pnas.1422089112.

Pinter-Wollman, Noa, Ashwin Bala, et al. (July 1, 2013). “Harvester Ants Use Interactions to Regulate Forager Activation and Availability”. In: Animal Behaviour 86.1, pp. 197–207. doi: 10.1016/j.anbehav.2013.05.012.

Pinter-Wollman, Noa, E. A. Hobson, et al. (Mar. 1, 2014). “The Dynamics of Animal Social Networks: Analytical, Conceptual, and Theoretical Advances”. In: Behavioral Ecology 25.2, pp. 242–255. doi: 10.1093/beheco/art047.

Pinter-Wollman, Noa, Roy Wollman, et al. (Nov. 7, 2011). “The Effect of Individual Variation on the Structure and Function of Interaction Networks in Harvester Ants”. In: Journal of The Royal Society Interface 8.64, pp. 1562–1573. doi: 10.1098/rsif.2011.0059.

Planckaert, Joffrey et al. (Oct. 30, 2019). “A Spatiotemporal Analysis of the Food Dissemination Process and the Trophallactic Network in the Ant Lasius Niger”. In: Scientific Reports 9.1 (1), pp. 1–11. doi: 10.1038/s41598-019-52019-6.

Psorakis, Ioannis et al. (Nov. 7, 2012). “Inferring Social Network Structure in Ecological Systems from Spatio-Temporal Data Streams”. In: Journal of the Royal Society, Interface 9.76, pp. 3055–3066. doi: 10.1098/rsif.2012.0223.

Quevillon, Lauren E. et al. (Aug. 24, 2015). “Social, Spatial, and Temporal Organization in a Complex Insect Society”. In: Scientific Reports 5, p. 13393. doi: 10.1038/srep13393.

Richardson, T.O. et al. (2020). Ant Behavioral Maturation Is Mediated by a Stochastic Transition between Two Fundamental States.

Robinson, Gene E. and Francis LW Ratnieks (1987). “Induction of Premature Honey Bee (Hymenoptera: Apidae) Flight by Juvenile Hormone Analogs Administered Orally or Topically”. In: Journal of Economic Entomology 80.4, pp. 784–787.

Rosenthal, Sara Brin et al. (Apr. 14, 2015). “Revealing the Hidden Networks of Interaction in Mobile Animal Groups Allows Prediction of Complex Behavioral Contagion”. In: Proceedings of the National Academy of Sciences 112.15, pp. 4690–4695. doi: 10.1073/pnas.1420068112.

Rueppell, Olav et al. (May 6, 2008). “Aging and Demographic Plasticity in Response to Experimental Age Structures in Honeybees (Apis Mellifera L)”. In: Behavioral Ecology and Sociobiology 62.10, p. 1621. doi: 10.1007/s00265-008-0591-7.

Salvatier, John, Thomas V. Wiecki, and Christopher Fonnesbeck (Apr. 6, 2016). “Probabilistic Program-ming in Python Using PyMC3”. In: PeerJ Computer Science 2, e55. doi: 10.7717/peerj-cs.55.

Scheiner, Ricarda (Aug. 1, 2012). “Birth Weight and Sucrose Responsiveness Predict Cognitive Skills of Honeybee Foragers”. In: Animal Behaviour 84.2, pp. 305–308. doi: 10.1016/j.anbehav.2012.05.011.

Scheiner, Ricarda, Robert E. Page, and Joachim Erber (2004). “Sucrose Responsiveness and Behavioral Plasticity in Honey Bees (Apis Mellifera)”. In: Apidologie 35.2, pp. 133–142.

Schneider, Stanley S. and Lee A. Lewis (Mar. 2004). “The Vibration Signal, Modulatory Communication and the Organization of Labor in Honey Bees, *Apis Mellifera*”. In: Apidologie 35.2, pp. 117–131. doi: 10.1051/apido:2004006.

Seeley, Thomas D. (Dec. 1, 1982). “Adaptive Significance of the Age Polyethism Schedule in Honeybee Colonies”. In: Behavioral Ecology and Sociobiology 11.4, pp. 287–293. doi: 10.1007/BF00299306.

Seeley, Thomas D. (Mar. 1, 1989). “Social Foraging in Honey Bees: How Nectar Foragers Assess Their Colony’s Nutritional Status”. In: Behavioral Ecology and Sociobiology 24.3, pp. 181–199. doi: 10.1007/BF00292101.

Seeley, Thomas D. (Dec. 1, 1992). “The Tremble Dance of the Honey Bee: Message and Meanings”. In: Behavioral Ecology and Sociobiology 31.6, pp. 375–383. doi: 10.1007/BF00170604.

Seeley, Thomas D. (1995). The Wisdom of the Hive: The Social Physiology of Honey Bee Colonies. Cambridge, Mass: Harvard University Press. 295 pp.

Seeley, Thomas D. et al. (Jan. 5, 2012). “Stop Signals Provide Cross Inhibition in Collective Decision-Making by Honeybee Swarms”. In: Science 335.6064, p. 108. doi: 10.1126/science.1210361.

Smith, M. L., M. M. Ostwald, and T. D. Seeley (Nov. 1, 2016). “Honey Bee Sociometry: Tracking Honey Bee Colonies and Their Nest Contents from Colony Founding until Death”. In: Insectes Sociaux 63.4, pp. 553–563. doi: 10.1007/s00040-016-0499-6.

Sosna, Matthew M. G. et al. (Oct. 8, 2019). “Individual and Collective Encoding of Risk in Animal Groups”. In: Proceedings of the National Academy of Sciences 116.41, pp. 20556–20561. doi: 10.1073/pnas.1905585116.

Strandburg-Peshkin, Ariana et al. (May 19, 2018). “Inferring Influence and Leadership in Moving Animal Groups”. In: Philosophical Transactions of the Royal Society B: Biological Sciences 373.1746, p. 20170006. doi: 10.1098/rstb.2017.0006.

Tofts, Chris and Nigel R. Franks (Oct. 1, 1992). “Doing the Right Thing: Ants, Honeybees and Naked Mole-Rats”. In: Trends in Ecology & Evolution 7.10, pp. 346–349. doi: 10.1016/0169-5347(92)90128-X.

Toth, Amy L. et al. (Dec. 2005). “Nutritional Status Influences Socially Regulated Foraging Ontogeny in Honey Bees”. In: The Journal of Experimental Biology 208 (Pt 24), pp. 4641–4649. doi: 10.1242/jeb.01956.

Traniello, James FA and Rebeca B. Rosengaus (1997). “Ecology, Evolution and Division of Labour in Social Insects”. In: Animal behaviour 53.1, pp. 209–213.

Traynor, Kirsten S., Yves Le Conte, and Robert E. Page (Jan. 1, 2015). “Age Matters: Pheromone Profiles of Larvae Differentially Influence Foraging Behaviour in the Honeybee, Apis Mellifera”. In: Animal Behaviour 99, pp. 1–8. doi: 10.1016/j.anbehav.2014.10.009.

Van der Walt, Stéfan et al. (June 19, 2014). “Scikit-Image: Image Processing in Python”. In: PeerJ 2, e453. doi: 10.7717/peerj.453.

Von Luxburg, Ulrike (Dec. 1, 2007). “A Tutorial on Spectral Clustering”. In: Statistics and Computing 17.4, pp. 395–416. doi: 10.1007/s11222-007-9033-z.

Wang, Ying, Osman Kaftanoglu, Colin S. Brent, et al. (Apr. 1, 2016). “Starvation Stress during Larval Development Facilitates an Adaptive Response in Adult Worker Honey Bees (Apis Mellifera L.)” In: Journal of Experimental Biology 219.7, pp. 949–959. doi: 10.1242/jeb.130435.

Wang, Ying, Osman Kaftanoglu, Adam J. Siegel, et al. (Dec. 1, 2010). “Surgically Increased Ovarian Mass in the Honey Bee Confirms Link between Reproductive Physiology and Worker Behavior”. In: Journal of Insect Physiology 56.12, pp. 1816–1824. doi: 10.1016/j.jinsphys.2010.07.013.

Wang, Ying, Sarah D. Kocher, et al. (Jan. 1, 2012). “Regulation of Behaviorally Associated Gene Networks in Worker Honey Bee Ovaries”. In: Journal of Experimental Biology 215.1, pp. 124–134. doi: 10.1242/jeb.060889.

Ward, Joe H. Jr. (1963). “Hierarchical Grouping to Optimize an Objective Function”. In: Journal of the American Statistical Association 58.301, pp. 236–244. doi: 10.1080/01621459.1963.10500845.

Wario, Fernando, Benjamin Wild, Margaret J. Couvillon, et al. (Sept. 25, 2015). “Automatic Methods for Long-Term Tracking and the Detection and Decoding of Communication Dances in Honeybees”. In: Frontiers in Ecology and Evolution 3. doi: 10.3389/fevo.2015.00103.

Wario, Fernando, Benjamin Wild, Raúl Rojas, et al. (2017). “Automatic Detection and Decoding of Honey Bee Waggle Dances”. In: PloS one 12.12, e0188626.

Wild, Benjamin, Leon Sixt, and Tim Landgraf (Feb. 13, 2018). Automatic Localization and Decoding of Honeybee Markers Using Deep Convolutional Neural Networks.

